# AI-guided discovery for low-resource peptide engineering using evolutionary scale modeling

**DOI:** 10.64898/2026.06.25.734678

**Authors:** Leo Andrekson, Robin Rydbergh, Rocío Mercado, Michaela Wenzel

## Abstract

Data scarcity is a major bottleneck in artificial intelligence (AI)-guided peptide engineering, where experimental training sets often contain only tens to a few hundred sequences, and it is difficult to judge in advance whether a protein language model (PLM) will effectively guide iterative discovery before committing to wet-lab work. Here we present SCARSE, a machine learning framework that couples ESM2 protein language model embeddings with Gaussian process regression. Benchmarked across 24 diverse peptide and small-protein datasets spanning substitution and indel variants, antimicrobial peptides, and hemolytic peptides, SCARSE significantly outperformed hand-engineered descriptor baselines on substitution and indel tasks in low-data regimes (20–500 training samples) and performed comparably on short non-mutant peptides. In active learning simulations it consistently prioritized high-performing candidates, discovering up to 7-fold more top-tier peptides than random sampling. Cross-validation (CV) R² correlated strongly with final active learning performance (Pearson r = 0.80–0.84), and evaluating on as few as 50 labeled peptides reduced CV R² variance enough for it to serve as a practical, data-efficient criterion for assessing model suitability before iterative screening. SCARSE thus provides an early diagnostic indicator for active learning success and is freely available as an open-source Python package (pip) and through an interactive Hugging Face Spaces web interface.

## Introduction

The rapid evolution of protein language models (PLMs) has transformed sequence-based prediction in computational biology, enabling the extraction of rich biological insights directly from amino acid sequences. Unlike traditional machine learning (ML) approaches that depend on manually engineered sequence descriptors, PLMs leverage self-supervised pre-training on vast unlabeled protein datasets to learn intrinsic sequence representations. This paradigm mirrors the revolution in natural language processing, where large transformer-based models, trained on massive text corpora, achieve state-of-the-art performance across diverse tasks through unsupervised pre-training followed by task-specific fine-tuning ^1,2^.

In biological systems, peptides constitute fundamental functional units with roles in immune signaling, enzymatic activity, and therapeutic applications. However, many sequence-to-function prediction problems in peptide engineering suffer from data scarcity due to experimental characterization being costly and time-consuming, leading to datasets that often contain only tens to a few hundred labeled examples ^3^. Standard PLMs, trained on millions of sequences, may not transfer effectively to these low-data regimes without appropriate adaptation strategies ^4^.

In active learning-driven peptide engineering, PLMs are increasingly used to provide pretrained representations that support iterative experimental design in data-limited settings (Fig. 1a). In this setting, PLMs provide rich sequence embeddings that can be coupled with lightweight supervised models to prioritize candidates for synthesis and testing, enabling efficient exploration of sequence space and enrichment of high-performing peptides over successive rounds. This approach has shown promise across tasks, where each experimental cycle benefits from model-informed selection rather than random screening.

**Fig. 1.**
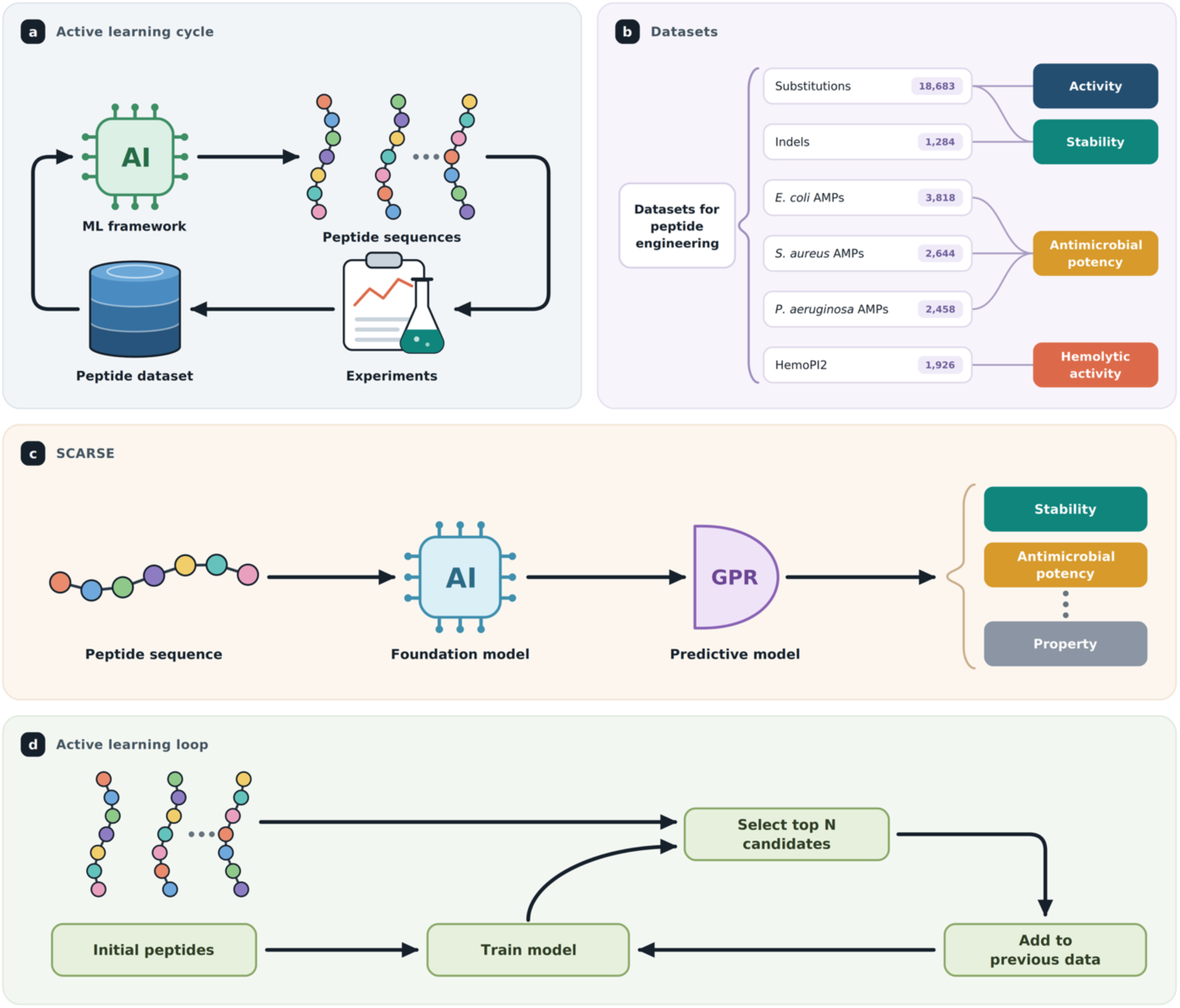
AI-infused peptide engineering concepts and the developed ML framework. (a) AI-infused peptide engineering workflow. The process starts with an initial dataset containing peptides with measured performance. This dataset is fed as training data to the ML framework, which then ranks the performance of a set of candidate peptides. Top N ranked peptides are selected for wet lab screening and added to the original dataset once screened. The cycle is repeated until a candidate with sufficient performance is found. (b) Datasets used to evaluate the ML framework, consisting of substitutions, indels, hemolytic peptides (HemoPI2), and antimicrobial peptides (AMPs) with potency measured against *Escherichia coli* (*E. coli*), *Staphylococcus aureus* (*S. aureus*), and *Pseudomonas aeruginosa* (*P. aeruginosa*). These datasets capture endpoints such as activity, stability, antimicrobial potency, and hemolytic activity. (c) Overall model architecture and data-processing pipeline of SCARSE. Raw peptide amino acid sequences are tokenized and embedded using the pretrained foundation model ESM2 650M before being passed to the predictive model, Gaussian process regressor (GPR), for supervised prediction of peptide properties. (d) Iterative active learning workflow simulation mimicking experimental discovery. The process begins with 20 randomly selected peptides for initial training. The trained model scores the remaining candidates, and the top-predicted peptides are iteratively added in batches of 20 over 10 rounds of candidate selection. This sequential retraining strategy emulates an active learning approach for data-efficient peptide optimization.

One approach leveraging a PLM to enable high-performance design of proteins with minimal experimental labels is eUniRep ^5^. Central to this approach is the use of a foundation model, UniRep, which is globally pre-trained on over 20 million amino acid sequences to distill fundamental biophysical and evolutionary features into a compact latent representation ^6^. This model is subsequently fine-tuned on sequences identified using JackHMMER that are similar to the target protein family to create eUniRep, a representation that captures both global protein attributes and local family-specific features ^5,7^. This approach showed that as few as 24 functionally characterized protein mutants could suffice to discover high-performing proteins, demonstrating potential benefits from integrating PLMs in the optimization workflow. One of the main differences between eUniRep and our ML framework is that our method does not fine-tune the foundation model and does not perform multiple sequence alignment (MSA), thereby reducing computational overhead. Eliminating the need for MSA is also advantageous when working with short amino acid sequences such as peptides, where alignment-based methods often perform suboptimally due to limited sequence length and reduced evolutionary signal, which can hinder reliable identification of homologous sequences.

Recent research has identified strategies that combine foundation models with few-shot learning frameworks to improve predictive accuracy of these models ^8^. For instance, the few-shot learning for protein fitness prediction (FSFP) methodology applied to ESM-1v increased the positive hit-rate compared to baseline when optimizing Phi29 DNA polymerase through single-site mutants ^8^.

Surveys provide broader context on the development and evaluation of PLMs, going through their use in tasks ranging from structural prediction to functional annotation and generation tasks ^9,10^. These reviews highlight both foundational architectures and the potential for integrating structure-aware features into PLMs for enhanced downstream prediction. This is also supported by the ProteinGYM benchmark, where methods using both sequence and structure information have shown great performance, such as ProSST and SaProt ^11–13^. However, a drawback of structure-based methods is the requirement of having the structures of all sequences. These need to either be experimentally acquired or produced using other ML methods, such as AlphaFold, which can be costly and time-consuming ^14^. An alternative approach involves methods that integrate MSA, which have demonstrated superior performance on the ProteinGym benchmark. However, it is also important to consider the drawbacks of incorporating MSA, particularly the substantial computational cost associated with generating them and the large variability in performance across different tasks ^10^. This highlights the computational benefit of using a simpler approach only using the sequence information as input, such as ESM2 ^15^.

Representation learning has also been applied directly to peptides, where models predict properties such as antibacterial activity, hemolysis, solubility, and antifouling properties. Bringing the PLM paradigm down to the peptide scale, PeptideBERT, built on the pretrained ProtBERT transformer, predicts these properties directly from sequence and reaches state-of-the-art hemolysis prediction, exploiting the fact that representing protein sequences as text enables sequence-to-property prediction without relying on explicit structural data ^16,17^. Arguing that sequence alone discards structural information, Multi-Peptide combines PeptideBERT with a graph neural network encoder and a contrastive loss framework to align sequence- and structure-based embeddings in a shared latent space, reaching 88% accuracy on hemolysis using structures generated with AlphaFold, though it failed to improve over the sequence-only model on antifouling ^14,18^. Others question the amino acid token itself, since most predictors are incompatible with non-natural features such as modified side chains or staples and rely on human-crafted amino acid features. PepMNet instead uses a hierarchical graph model that learns from an atom-level graph before forming amino acid and molecular representations, and PepFoundry encodes peptides beyond canonical amino acids and linear topologies as SMILES strings in the CHUCKLES format, with atomic-level representations consistently outperforming sequence-level ones ^19–23^. A common limitation across these approaches, however, is that they are trained and benchmarked on relatively large datasets, leaving their performance in the low-data regimes typically encountered in practical peptide engineering largely unexamined. This matters because, in practice, it is often unclear prior to committing resources to an active learning campaign whether the available data are sufficient to support effective model generalization and consequently, what level of performance or improvement can realistically be expected from integrating PLMs with iterative experimental workflows. This gap is partially compounded by the limited understanding of how foundation model representations perform when coupled with downstream regression and classification architectures under low-data and short-sequence conditions, where most peptide engineering problems reside. While benchmark studies and workflow simulations can address these questions by evaluating model behavior across diverse datasets and emulating iterative discovery processes, they do not directly provide a practical criterion for deciding, in advance, whether a given dataset and modeling strategy will succeed in an active learning setting. Together, these challenges motivate the need for a simple, data-efficient method that can reliably predict active learning performance prior to experimental commitment, enabling informed decisions about when and how to deploy PLM-guided peptide engineering workflows.

To address these challenges, this study introduces SCARSE: Small-sample Cross-validation-Anchored Regression Solution for low resource peptide Engineering. SCARSE is an AI-based framework that can be trained using very few samples and can predict peptide properties based on sequences only, making for a fast and high-performing approach for peptide engineering. We assess performance across varying datasets, training sizes, and simulated active learning scenarios that mirror iterative peptide optimization. We also present a method that enables early estimation of final active learning performance, allowing evaluation before full simulation or experimental completion. Finally, SCARSE is deployed in a user-friendly interface to facilitate adoption of peptide activity prediction and optimization into peptide engineering pipelines.

## Methods

### AI use declaration

This work made use of large language model (LLM)-based artificial intelligence tools, more specifically ChatGPT-4o, Claude Sonnet 4.6 and Opus 4.8, and Gemini 3.1 Flash-Lite, 3.5 Flash, and 3.1 Pro, for support in text editing and coding. These tools were used to assist with improving clarity, grammar, and overall readability of the written content, as well as to provide suggestions and guidance in the development and refinement of code.

### Data curation

To evaluate the generalizability of SCARSE, multiple types of peptide datasets were investigated, including substitutions, indels, antimicrobial peptides (AMPs), and hemolytic peptides (Fig. 1b). In this work, we collected data from ProteinGYM, one of the largest publicly available benchmarks for deep mutational scanning (DMS) assays of protein and peptide mutational effects, containing many datasets of varying sequence lengths ^11^. We selected 10 datasets (Table S1, Section S1) containing one and two-point substitution mutants with sequence lengths ranging from 39 up to 87 amino acids (aa). These datasets cover mammalian, archaeal, bacterial, and viral sources, and consist of a total of 18,683 characterized sequences with the endpoint defined as stability or activity. These were picked for their relatively short sequence lengths compared to other datasets in ProteinGYM. We also collected 10 indel datasets (Table S2, Section S1). These datasets cover mammalian, archaeal, bacterial, and viral sources, and consist of a total of 1,284 characterized sequences of length ranging from 39 to 71 aa with the endpoint defined as stability. It is worth noting that the number of sequences in each of the indel datasets is substantially fewer than for substitutions, putting constraints on computational evaluations. The endpoint distributions of the substitution and indel datasets are shown in Fig. S1 and Fig. S2 (Section S1).

Finally, to evaluate model performance on very short peptides we included three AMP potency datasets, each collecting the potency of experimentally validated AMPs against a single target organism, namely *Escherichia coli* (3,818 sequences), *Staphylococcus aureus* (2,644 sequences), and *Pseudomonas aeruginosa* (2,458 sequences), with lengths ranging from 6 to 40 aa ^24^. The endpoint for these three datasets is pMIC, defined as −log₁₀(MIC), the negative base-10 logarithm of the minimal inhibitory concentration (MIC). In addition, we included a hemolytic-activity dataset, HemoPI2, comprising 1,926 sequences of length 6 to 39 aa, with the endpoint defined as pHC₅₀ = −log₁₀(HC₅₀), the negative base-10 logarithm of the concentration causing 50% hemolysis (HC₅₀) ^25^ (Table S3, Section S1). All datasets used in this study exclusively contained the 20 canonical L-amino acids, represented by their standard IUPAC one-letter codes. Levenshtein distance was used for filtering out sequences that have a distance of 2 or less between each other ^26^. This approach was applied to the AMP and HemoPI2 datasets to increase the dissimilarity of sequences in order to reflect a scenario where sequences are further from each other than two-point mutations. The endpoint distributions of these peptide datasets are shown in Fig. S3 (Section S1).

### Model framework

The overall framework of SCARSE consists of two main components, the input representation generated by ESM2, which produces fixed-length peptide sequence embeddings, and an ML model (a supervised regressor) that maps these embeddings to quantitative experimental outcomes, such as stability, activity, or hemolytic activity, for instance (Fig. 1c). We also deployed three baseline input representations for performance comparison.

### Input representation

For the input representation of SCARSE, we employed the ESM2 family of protein language models. More specifically, ESM2 650M parameters was used due to its strong balance between representational power and computational efficiency ^15^. A comparison across ESM2 model sizes (8M, 35M, 150M, and 650M parameters) showed that predictive performance gains diminished with size, with little improvement from 150M to 650M, so we used the largest model tested, 650M (Fig. S4, Section S2). Sequence embeddings were extracted by mean-pooling the final hidden layer representations.

The baseline comprised three sequence representations that do not rely on a protein language model, referred to as Physchem, Morgan, and N-gram. Descriptors calculated for the Physchem representation consist of all global descriptors from the modlAMP Python package ^27^.

Additionally, we also included amino acid percentage composition, the percentage of polar (S, T, N, H, Q, G, E, D, K, R), apolar (A, L, V, I, M, W, Y, F, P, C), charged (E, D, K, R), and aromatic (W, Y, F) amino acids, and finally hydrophobic moment and discrimination factor inspired by HeliQuest ^28,29^. This selection gives a simple yet robust set of features capable of peptide property prediction. The Morgan representation encodes each peptide as a Morgan (circular) fingerprint computed from its molecular structure, and the N-gram representation encodes each peptide as term frequency-inverse document frequency (TF-IDF) weighted character n-grams of order 1–3, with the vocabulary fit on the training sequences only. Together, these three baselines span physicochemical, structural, and compositional descriptor families. The settings for all three baseline representations are summarized in Section S3, Table S5.

Both ESM2 and the baseline representations were normalized using the StandardScaler function from scikit-learn ^30^. Within each cross-validation split the scaler was fitted on the training fold and then applied to transform the held-out validation fold, and for the final evaluation it was fitted on the full training set and applied to transform the test set. Fitting the scaler only on training data ensures that no validation or test statistics leak into the features.

### Predictive models

To assess the predictive utility of the input representations, we applied the Gaussian process regressor (GPR) for all prediction tasks ^31^. GPR was selected for its robustness in small-sample settings.

### Hyperparameter tuning

For hyperparameter tuning we leveraged the Optuna optimization framework ^32^. For each training size and random seed, Optuna was instructed to perform 100 trials to search the hyperparameter space (Table S4, Section S3). An optimum was achieved by minimizing the average mean squared error (MSE) loss across a 10-fold cross validation (CV). Splits were performed randomly for CV. SCARSE inherently includes hyperparameter tuning as part of its workflow.

### Evaluation strategy for benchmarking

Each training iteration involved selecting a random seed and a training set size, followed by partitioning the data into 10 CV folds. Model performance was then evaluated on a held-out test set where the benchmarking performance was reported using the R² and Spearman ρ metrics. For the substitution, AMPs, and HemoPI2 datasets, the test set consisted of 500 data points. For the indel datasets, the test set size was reduced to 20 data points.

To account for variability and improve robustness, 10 random seeds were used for each training set size. The following training sizes were evaluated for the substitution, AMPs, and HemoPI2 datasets: [20, 50, 75, 100, 200, 350, 500]. For the indel datasets, the evaluated training sizes were [20, 40, 60, 80].

Two distinct strategies were employed to partition the data into training and test sets. For the substitution and indel datasets, samples were assigned using random splitting. In contrast, for the AMPs and HemoPI2 datasets, a similarity-aware splitting approach was implemented to minimize sequence similarity between training and test sets while preserving both endpoint and sequence length distributions.

This procedure was conducted on a per-sequence length basis. First, the test set was constructed by randomly selecting an initial sequence for each sequence length. Subsequently, additional sequences of the same length were iteratively included based on minimal Levenshtein distance to the initially selected sequence, until the number of samples matched the proportion dictated by the overall sequence length distribution. For example, if the target test set size was 500 sequences and 10% of the dataset consisted of sequences of 10 aa length, then 50 such sequences were included in the test set: one randomly selected sequence, followed by the 49 most similar sequences (i.e., those with the smallest Levenshtein distance).

The training set was then constructed by selecting, for each sequence length, those sequences that were maximally dissimilar (i.e., with the largest Levenshtein distance) from the corresponding test set seed sequence, again respecting the underlying sequence length distribution.

This strategy ensures that the training and test sets are well separated in sequence space, as defined by Levenshtein distance, while maintaining key distributional properties required for robust model generalization. The rationale is to approximate a realistic prediction scenario, in which the model is evaluated on sequences that are dissimilar from those encountered during training.

Both ESM2 and the baseline descriptors were included in the benchmark of each dataset. Statistically significant differences between SCARSE and each baseline representation were assessed using the Tukey HSD posthoc test, which has previously been recommended for small-molecule drug discovery ^33^.

For the SDA BACSU dataset the Physchem baseline gave unstable predictions with strongly negative R² for a few random seeds at training sizes 50, 75 and 100. To prevent these outliers from distorting the figures we excluded random seeds with a test R² below −10 from that dataset’s Physchem results when visualizing the benchmarking results. The statistical tests were carried out on the full unfiltered data (see Section S4).

### Active learning simulations

To evaluate SCARSE in a laboratory-relevant setting, we simulated an iterative peptide discovery workflow mimicking active learning (Fig. 1d). For each experiment, the workflow was initialized with 20 randomly selected peptides, and this initialization was repeated across 10 different random seeds, corresponding to 10 distinct starting sets of 20 peptides. The model was trained on the initial set and used to predict the remaining candidates. The top 20 peptides with the highest predicted values were then added to the training set, after which the model was retrained. This iterative process was repeated until 200 samples were collected (10 rounds total). This greedy top-k selection was compared against a mixed acquisition strategy, in which each round added 70% of its batch from the highest-predicted candidates and 30% from the lowest-predicted candidates, introducing a degree of exploration alongside exploitation. Model-guided selection was compared to random sampling and the three descriptor baselines (Physchem, Morgan, and N-gram), and performance was recorded after each round. Experiments were conducted across all datasets, except indel-based datasets.

To quantify the ability of the models to identify high-performing peptides, we used the “Top 10% peptides selected (%)” metric. This metric measures the percentage of the true top 10% highest-performing peptides in the full dataset that had been identified after screening a given number of peptides. Evaluation using this metric was applied to the substitution, AMPs, and HemoPI2 datasets. For all datasets a higher endpoint value was treated as more desirable, with one exception. For HemoPI2 the endpoint was pHC₅₀ = −log₁₀(HC₅₀) where a lower value corresponded to a higher tolerated concentration and thus a less hemolytic and more desirable peptide. HemoPI2 was therefore optimized in the opposite direction, so that the true top 10% and every selection step targeted the lowest pHC₅₀ values. In addition, we assessed the final active learning enrichment performance of SCARSE by calculating the ratio between the final top 10% selection performance achieved by SCARSE after 200 screened peptides and that achieved by random sampling. Expressing performance as a multiple of random screening places all datasets on a common scale, since the raw top 10% selection value depends on the pool size and on how many of the true top 10% a random draw would recover for each dataset. A value of 1 corresponds to random-sampling performance and values above 1 indicate enrichment over random. Finally, we examined how this enrichment performance related to model predictive quality by comparing it to CV R² scores obtained from models trained on random subsets of varying sizes from each dataset.

## Results

### Benchmark performance

Fig. 2a reports the average performance of SCARSE next to the baseline representations across all substitution datasets, together with the best-performing (Fig. 2b) and the worst-performing substitution datasets (Fig. 2c). We note that relatively high R^2^ values are achievable even with low amounts of training data, but also that the performance of the models is highly dataset-dependent. This dataset dependence is evident in the learning curves (Fig. S5, Section S4): for some datasets an R² value of 0.72 is reached at the smallest training size of 20 (SDA BACSU), for others an R² value of 0.53 is achievable at a training size of 100 (SOX30 HUMAN), while still others remain below an R² value of 0.29 even at the maximum training set size of 500 (ENVZ ECOLI). However, SCARSE is statistically significantly better than the baseline representations for most substitution datasets and training sizes (Fig. S5k-m, Section S4). The Spearman ρ rank correlation for the substitution datasets is shown in Fig. S6 (Section S4).

**Fig. 2.**
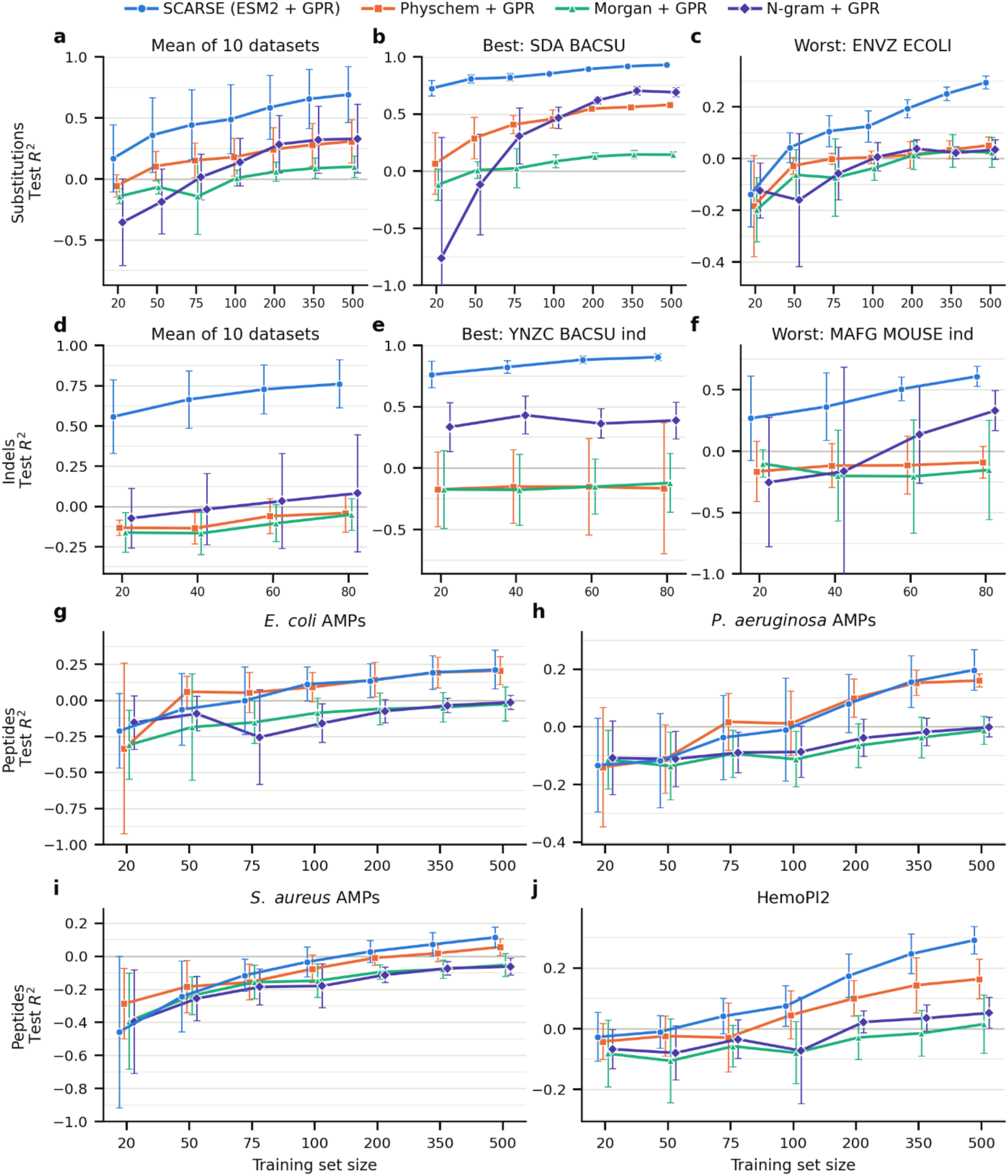
Benchmarking results. Test R² is shown for SCARSE (ESM2 + GPR) and the three descriptor baselines (Physchem, Morgan, and N-gram, each combined with GPR) as a function of training set size. (a) Mean across all 10 substitution datasets. (b) Best performing substitution dataset (SDA BACSU). (c) Worst performing substitution dataset (ENVZ ECOLI). (d) Mean across all 10 indel datasets. (e) Best performing indel dataset (YNZC BACSU ind). (f) Worst performing indel dataset (MAFG MOUSE ind). (g–j) The antimicrobial and hemolytic peptide datasets: (g) *E. coli* AMPs, (h) HemoPI2, (i) *P. aeruginosa* AMPs, and (j) *S. aureus* AMPs. Values are the mean R² across 10 random seeds, and for the aggregate panels (a, d) additionally averaged across datasets; error bars show the standard deviation (std) across seeds, or across datasets for the aggregate panels. Per-dataset benchmarking curves and Tukey HSD significance testing against each baseline are provided in Section S4.

On the indel datasets, SCARSE achieved a mean R^2^ value of 0.55 already at the smallest training set size of 20, rising to a R^2^ value of 0.76 at a training size of 80 (Fig. 2d). This is in contrast to the baseline representations, which yielded a mean R² value below or slightly above 0 across training sizes, performing statistically significantly worse than SCARSE across most datasets and training sizes. For indel-based datasets we also observed a clear performance dependency on the dataset (Fig. 2e,f). Per-dataset indel benchmarking curves are shown in Fig. S8 (Section S4).

Fig. 2g–j illustrates the benchmarking performance on the antimicrobial and hemolytic peptide datasets. On the three AMP potency datasets, SCARSE performed on par with the strongest baseline, Physchem, with both clearly ahead of the Morgan and N-gram representations, whereas on HemoPI2 SCARSE reached the highest R² of all representations at the larger training sizes.

Overall predictive performance on these short, compositionally diverse peptides was modest, with statistically significant gains for SCARSE emerging mainly at larger training sizes and against the Morgan and N-gram baselines (Section S4). Per-dataset peptide results, along with Tukey HSD post-hoc statistical tests, are shown in Fig. S9, and the corresponding Spearman ρ rank correlation in Fig. S10 (Section S4).

When applying SCARSE for predicting properties of unseen peptides in a peptide engineering scenario, its predictive performance on the sequences it predicts to have the most desirable quality, meaning its most extreme predictions, would be of particular interest. For the substitution, AMPs, and HemoPI2 datasets, we quantified the top 20 accuracy at different training sizes (Fig. S7, S11, Section S4). On average the models outperform random sampling across training sizes and SCARSE outperforms the baselines in many of the evaluated datasets. We note that random sampling would be equivalent to a top 20 accuracy of 4% on average. It is, however, worth mentioning that the data dependency seen for the substitution, AMPs, and HemoPI2 datasets in Fig. 2 also carries over to the selection of high-performing peptides, quantified per dataset by the Top 20 accuracy in Fig. S7 for the substitution datasets and Fig. S11 for the AMPs and HemoPI2 datasets (Section S4).

### Active learning simulation performance

Indications of high selection performance in Fig. S7 and S11 (Section S4) hinted at the opportunity of using SCARSE in a lab-like setting, paving the way for further investigations. Looking at the overall iterative peptide selection performance of SCARSE (the per-round and cumulative metrics are defined in Section S5; Fig. S12, S13, Section S6), we note how the average peptide selected by the model indeed tends to have enhanced performance compared to random sampling. Although the overall cumulative performance increased more gradually as additional peptides were screened (Fig. S13, Section S6), analysis of how SCARSE selected the top available peptide candidates during the active learning simulations revealed a different trend. Specifically, the model showed an almost linear increase in the number of high-performing peptides identified as the number of peptides screened grows. In addition, SCARSE consistently outperformed random sampling across the simulations, demonstrating its ability to efficiently prioritize high-performing peptide candidates (Fig. 3a-g).

**Fig. 3.**
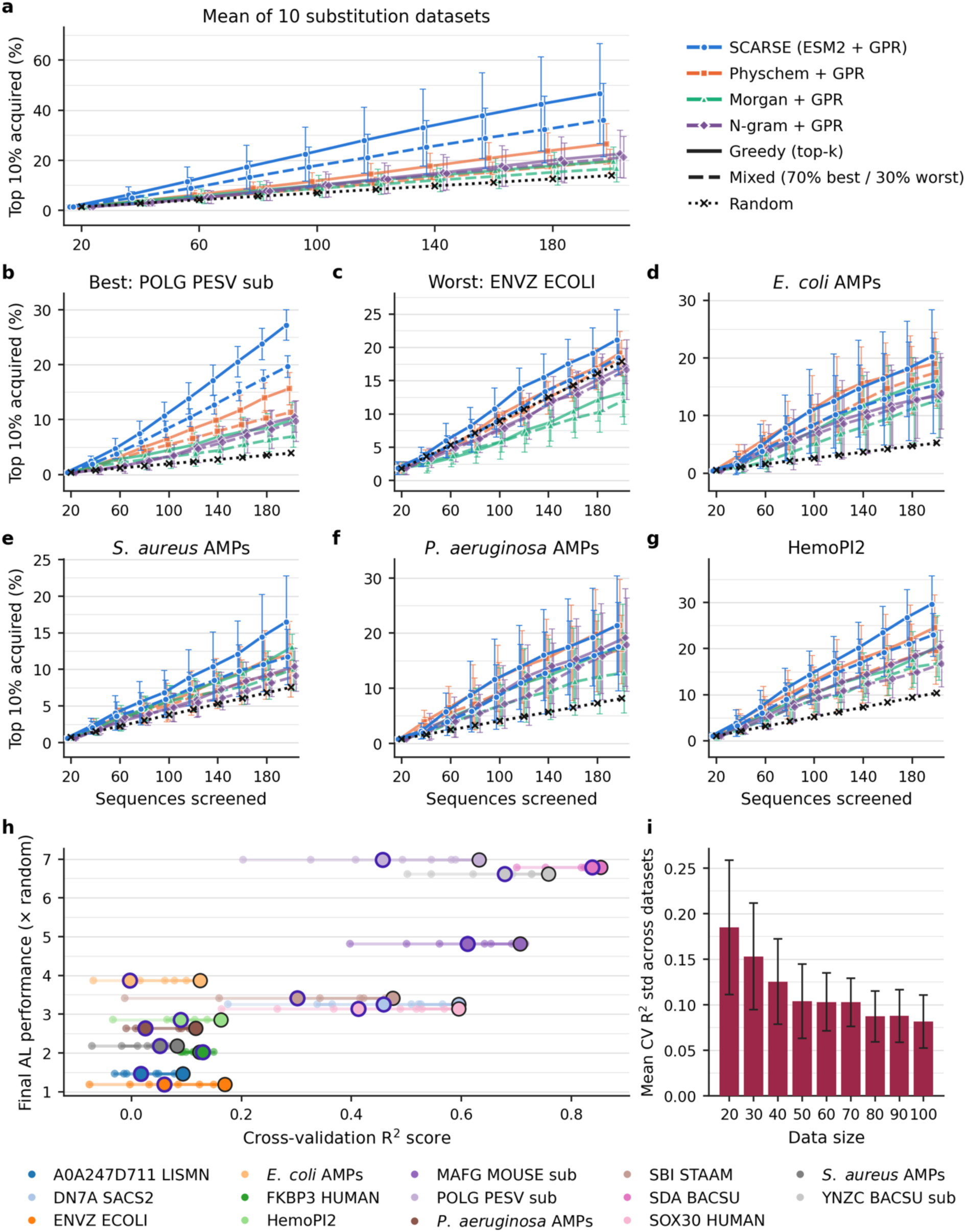
Active learning simulation results. (a) Percentage of the actual top 10% of peptides acquired by SCARSE and the three baselines up to each selection round, aggregated across the 10 substitution datasets. Solid lines show greedy top-k selection, dashed lines the mixed (70% best / 30% worst) strategy, and the dotted line random sampling. Values represent the mean across datasets and 10 random seeds, with the std across datasets. (b) Best-performing substitution dataset (POLG PESV sub). (c) Worst-performing substitution dataset (ENVZ ECOLI). (d– g) The antimicrobial and hemolytic peptide datasets: (d) *E. coli* AMPs, (e) *S. aureus* AMPs, (f) *P. aeruginosa* AMPs, and (g) HemoPI2. (h) Correlation between the final active learning (AL) performance (after 10 rounds and 200 screened peptides) and CV R² score when using SCARSE. The y-axis shows the final AL top 10% acquisition performance under the greedy top-k strategy from the workflow simulations (Fig. S14, Section S6) divided by the performance of random sampling, so a value of 1 equals random sampling. The x-axis shows the CV R² obtained from random subsets of size 20–100, shown as a color gradient from 20 (most transparent) to 100 (least transparent, black outline) for each dataset, where the 50 data size mark has a dark purple outline. (i) Mean std of the CV R² scores across datasets at different random-subset sizes, with std error bars across datasets. For each dataset and data size, 10 random seeds were used. The mean standard deviation of the final AL performance across datasets when using SCARSE is 0.53 under the greedy strategy (0.46 under the mixed strategy).

Towards the end of the workflow simulations (Fig. S14, Section S6), SCARSE discovers up to 7 times more high-value peptides compared to random sampling, further validating the use of SCARSE for peptide engineering. We further note that SCARSE on average had higher performance than the baselines on the substitution datasets (Fig. 3a), while also being a top-performer on the AMPs and HemoPI2 datasets (Fig. 3d-g). However, the issue remains that active learning performance heavily depends on the data at hand. The final active learning performance also varied across the 10 random seeds and this spread was widest for the antimicrobial and hemolytic datasets and grew as more peptides were screened (Fig. S15a–c, Section S6). The spread appeared under both the greedy and the mixed strategy and was only weakly to moderately related to the composition of the initial training set, whether measured by the spread or the best value of the initial labels or by the sequence-length range of the initial set (Fig. S15d–f). Despite this spread, every individual run still outperformed random sampling.

Getting an approximation of active learning workflow performance from a small amount of initial data would aid researchers in deciding whether SCARSE is a suitable framework for their data or if investigations into other candidate selection strategies may be warranted. Hence, we investigated whether the CV R^2^ score based on small random subsets of the datasets correlates to the final active learning performance (after 10 rounds and 200 screened peptides) of the active learning workflow simulations (Fig. 3h, Fig. S16, Section S6). The strength of this approach is that the CV R^2^ score can always be calculated, providing an estimate of active learning performance only based on currently existing data. Fig. 3h, i shows that there is a clear correlation between final active learning performance and CV R^2^ score, with a Pearson correlation coefficient ranging from 0.80 to 0.84 depending on the size of the random subsets of data, showing that this is indeed feasible.

## Discussion

### Benchmarking

Fig. 2a,d show that SCARSE consistently outperforms the baseline representations. We attribute this improvement to the use of ESM2, which is able to distinguish between highly similar sequences and encode them into generalizable embeddings. This could then be leveraged by the GPR model to make predictions. In contrast, the relatively simple descriptor baselines may not be complex enough to capture the variation from these mutants, resulting in worse generalizability of the GPR.

From observing the benchmarking performance in Fig. 2g–j, we note that the AMPs and HemoPI2 datasets demonstrate limited generalization performance on unseen data, requiring up to 500 characterized peptides to reach only modest R² values. However, even though the overall performance is not ideal, the accumulation of high-performing peptides seems to be much greater than one might expect from looking at the overall R^2^ alone (Fig. S9, S11, Section S4). This motivates the analysis of high-performing peptides as a complement to the overall R^2^ benchmarking.

### Active learning simulations

Both in the benchmarks and active learning simulations, we observe that the performance heavily depends on the dataset. SCARSE tends to outperform the baselines on datasets consisting of one and two-point mutants while performance similar to the strongest baseline is achieved on shorter and more compositionally diverse sequences, such as on the AMP datasets. We hypothesize that ESM2 captures the effect of single and double amino acid mutations more effectively than the physicochemical, structural, and compositional descriptors of the three baselines. A likely reason is that ESM2 is a context-dependent sequence model trained on large-scale protein data whereas the baseline descriptors summarize a sequence with fixed features that do not model how specific residues interact. Substitution and indel tasks reward exactly this sensitivity to individual sequence changes which may explain the improved performance. Confirming this would require directly probing the learned representations which is beyond the scope of this work. In contrast, on the short and compositionally diverse peptide datasets this advantage largely disappears and the strongest hand-engineered baseline, the physicochemical descriptors, becomes competitive with SCARSE. However, we do note that the selection performance of high-performing peptides during active learning for datasets A0A247D711 LISMN and ENVZ ECOLI (Fig. S14a, c, Section S6) is similar between SCARSE and the Physchem baseline, even though they are both substitution-based datasets. One distinction between these datasets and the rest is that the endpoint is activity. One might assume this is the cause of the deviation, but upon further inspection we observe that the endpoint distribution of dataset A0A247D711 LISMN is very similar to that of the stability endpoint dataset DN7A SACS2 (Fig. S1, Section S1), but SCARSE achieved much greater performance on the DN7A SACS2 dataset compared to the baselines (Fig. S14a, b, Section S6). This indicates that the reason for the deviation is likely not only caused by the endpoint distribution. Further investigation into the effects of endpoint, sequence length, and sequence diversity would be required to determine the extent to which these factors influence the observed outcomes.

One common theme of the active learning simulations across datasets is that the selectivity for high-performing peptides remains largely unchanged during the simulated accumulation of 200 peptides (Fig. S14, Section S6). It appears that the added knowledge gained from each selection round adding more peptides to the training data counteracts the reducing number of high-performing peptides available at each round, resulting in an almost linear increase of selected high-performing peptides in relation to the number of peptides screened. These results suggest that AI-infused peptide engineering using SCARSE is just as advantageous early in the process as in later stages where 200 peptides have been screened, indicating that SCARSE can be beneficially implemented at any stage of the peptide engineering process.

The averaged curves also conceal a spread across random seeds that is largest for the antimicrobial and hemolytic peptide datasets and widens as more peptides are screened (Fig. S15a–c, Section S6). This spread is present under both the greedy and the mixed strategy (Fig. 3 and Section S6), indicating that it is not a consequence of pure exploitation. It is also only weakly to moderately related to the composition of the initial 20-peptide training set, whether measured by the spread or the best value of the initial labels or by the sequence-length range of the initial set (Fig. S15d–f, Pearson r ≤ 0.30, Section S6). Since the initial set accounts for little of the spread, the variance more plausibly arises from the round-to-round accumulation of training data on these particular datasets, which remains to be confirmed directly. This further highlights the dataset dependence of using ML to guide peptide engineering.

The high correlation between CV R^2^ score and the final active learning performance observed in Fig. 3h indicates that calculating the CV R^2^ scores enables a quick and simple assessment of the suitability of a given dataset for active learning workflows using SCARSE. Even though correlation is high across data sizes, the standard deviation along the CV R^2^ score is large for very small data sizes, making it unreliable (Fig. 3i). We conclude that as little as 50 data points (Fig. 3h, Fig. S16, Section S6) can suffice to get a reasonably reliable indication of workflow performance from the CV R^2^ score.

We propose that users of SCARSE start by generating the CV R^2^ performance on their data and compare it to Fig. 3h and Fig. S16 (Section S6) to obtain an indication of how suitable SCARSE is for peptide candidate selection in an active learning workflow scenario for the dataset in question. To increase reliability of this estimate, we recommend use for cases where 50 or more peptides have been characterized. While starting with just 20 randomly selected peptides can suffice to achieve great performance (Fig. 3), reliable estimation of final active learning performance prior to completing all wet-lab experiments requires 50 or more characterized samples. However, even in situations where SCARSE displays poor modeling performance but no better methods are known for selecting peptide candidates, SCARSE would still be a viable option due to it performing better than random sampling for all tested datasets (Fig. S14, Section S6).

### Limitations

We also want to highlight the limitations of this work, primarily the limited number of peptides in the candidate pool that SCARSE selected from during the active learning simulations. In a lab-based peptide engineering pipeline, the set of possible peptide candidates that the ML framework predicts on may be much larger and may even change over selection rounds. The active learning simulations of this work provide an indication that AI-guided candidate selection outperforms random sampling, but the effects of larger and evolving peptide selection pools remain to be explored through *in vitro* AI-infused peptide engineering experimentation. We also want to emphasize that the active learning simulations implemented in this study relied on two selection strategies: a greedy top-k strategy, in which the model selects the top-ranked candidates at each iteration, and a mixed strategy that additionally samples a fraction of the lowest-predicted candidates to encourage exploration. The greedy strategy remained the strongest performer across all datasets (Fig. 3 and Section S6). Neither approach incorporates uncertainty-based acquisition functions commonly used in conventional active learning frameworks, and a systematic study of such acquisition functions, which explicitly balance exploration and exploitation, remains an avenue for future work. We further observed that the seed-to-seed spread was comparable between the greedy and mixed strategy on the substitution, antimicrobial and hemolytic peptide datasets (Fig. S14, Section S6). Whether uncertainty-aware acquisition functions would improve the reproducibility of individual runs on these datasets therefore remains an open question rather than an expected outcome.

This work has further limitations regarding the explored endpoints. While we have investigated different endpoints including activity, stability, antimicrobial potency, and hemolytic activity, many other peptide properties may be of interest. We acknowledge that the performance of SCARSE reported in this work may not translate to other peptide properties. To get a better understanding of the validity of SCARSE for other datasets, we encourage the user to calculate the CV R^2^ value and compare it to Fig. 3h and Fig. S16 (Section S6) in order to get an indication whether the developed ML framework is capable of generalizing to a point where active learning workflows reach a desirable expected outcome.

In this work, we only evaluated performance on datasets with a single endpoint. One can however imagine scenarios where the optimal peptide depends on multiple endpoints, e.g., pMIC and hemolysis for AMPs. In such cases, SCARSE can first be used to predict each endpoint, after which one defines a loss function for weighting these attributes. However, this approach introduces the challenge of determining the relative importance of each endpoint. We recommend that implementations of AI-driven peptide engineering carefully consider how these weights are assigned, as inappropriate weighting can significantly bias the active learning workflow and misrepresent the desired properties of the target peptide.

### Future research

#### Peptide candidate pool

A crucial component of any iterative active learning workflow in peptide engineering is the definition of the candidate space, the set of peptide sequences from which the model selects or ranks potential high-performing variants. The way this space is defined directly impacts both the efficiency of exploration and the diversity of discovered peptides. We hypothesize that investigation into candidate generation strategies could lead to better outcomes for peptide engineering. This is motivated by the relatively large standard deviation in terms of the final active learning performance of some of the datasets in Fig. S16 (Section S6), showing that the selection of peptides at the start of the active learning workflow may impact the outcome at the end of the active learning campaign.

One common strategy involves generating mutational libraries, such as all possible single-point substitutions or small combinatorial expansions around a known high-performing peptide. This approach ensures local exploration around experimentally validated scaffolds, allowing the model to infer how specific amino acid changes affect properties, such as activity, stability, or binding affinity. In the Results section we confirm that high performance is achievable through such approaches.

Alternatively, researchers may define a static candidate pool in advance, comprising either experimentally characterized sequences, literature-curated datasets, or *in silico*–generated peptides from generative models. In this mode, SCARSE functions as a ranking engine, scoring all candidates to prioritize the most promising ones for downstream testing. This setup is particularly useful in early-stage discovery when an external library, such as the Antimicrobial Peptide Database (APD) or a focused sequence repository is already available ^34^. Another benefit of maintaining a fixed candidate space is that model evaluation becomes highly reproducible.

More sophisticated workflows can have sets of candidates that update as new data is acquired. For instance, after each training round, top-performing sequences can serve as seeds for generating new candidates via mutation or generative sampling, gradually expanding the search space toward untested but high-potential regions.

In the present study, we simulate an active learning scenario by selecting top-ranked peptides from a fixed dataset at each iteration. However, the same framework could easily extend to adaptive mutation-based or generative candidate spaces, enabling integration with *in silico* peptide design or directed evolution experiments.

#### Filtering

Another promising addition to AI-infused peptide engineering pipelines are filtering steps, which enable the removal of peptide candidates based on a set of conditions. The filter would act as a sanity check to the sequences in the peptide candidate pool, making sure that sequences with unreasonable properties are not considered. This would however only be beneficial when optimizing for a specific group of peptides where condition boundaries can be clearly defined, such as sequence length or hydrophobicity. Filtering would need to be manually defined for the peptide set in question, necessitating custom solutions on a case-by-case basis. We propose that investigation into the viability of such filters in AI-infused peptide engineering can be beneficial and further improve the probability of discovering high-performing peptides.

## Conclusions

In this work, we systematically investigated how training set size impacts machine learning performance for sequence-based peptide property prediction and, importantly, how CV R² score can be used as a practical indicator of downstream success in active learning workflows. Across experiments, we found that models built on ESM2 650M embeddings achieved the strongest performance, with GPR as the downstream regression model, as assessed by benchmarking and active learning simulations. Notably, SCARSE maintained strong predictive power even with training sets as small as 20 samples, underscoring its utility in data-limited peptide engineering scenarios. SCARSE further demonstrated consistent enrichment of high-performing peptides, a key requirement for effective AI-guided optimization, and in simulated active learning workflows, model-guided selection in many cases substantially outperformed random sampling. Crucially, we showed that CV R² computed on as few as 50 labeled peptides can be sufficient to estimate the eventual final performance of active learning scenarios, providing researchers with a simple, data-efficient criterion for evaluating performance of a given modeling setup before committing to iterative experimental cycles. Finally, SCARSE is made available as both a pip package and via HuggingFace Spaces to support integration into low-resource peptide engineering pipelines.

## Author contributions

**LA:** Conceptualization, Investigation, Methodology, Software, Validation, Data Curation, Writing - Original Draft, Visualization. **RR:** Validation, Writing - Review & Editing. **RM:** Conceptualization, Investigation, Methodology, Resources, Writing - Original Draft, Writing - Review & Editing, Visualization, Supervision. **MW:** Conceptualization, Investigation, Writing - Review & Editing, Supervision, Resources, Funding acquisition, Project administration.

## Data availability

The ProteinGYM datasets supporting the conclusions of this article are available in the ProteinGYM repository, https://proteingym.org/download.

The AMP potency and HemoPI2 datasets supporting the conclusions of this article are available from their original sources. The AMP datasets, https://doi.org/10.1016/j.isci.2024.110718. The HemoPI2 dataset, https://doi.org/10.1038/s42003-025-07615-w.

Code for SCARSE can be accessed below:

- Project name: SCARSE
- Project pages:

`o` HuggingFace application: https://huggingface.co/spaces/SCARSE/scarse
`o` GitHub implementation: https://github.com/LeoAnd00/SCARSE
`o` Reproducibility code: https://github.com/LeoAnd00/SCARSE-repro
- Archived version: 10.5281/zenodo.23234000
- Operating system(s): Platform-independent
- Programming language: Python
- Other requirements: Python ≥3.12.10, pip
- License: MIT
- Any restrictions to use by non-academics: None

## Acknowledgements

This work was funded by the Swedish Research Council (grant number 2024-02040 to MW). RM acknowledges funding provided by the Wallenberg AI, Autonomous Systems, and Software Program (WASP), supported by the Knut and Alice Wallenberg Foundation. Funders had no role in study design, data collection and interpretation, or the decision to submit the work for publication.

Computational resources were provided by the National Academic Infrastructure for Supercomputing in Sweden (NAISS), funded by the Swedish Research Council. Computational resources were provided on the Berzelius system funded by the Knut and Alice Wallenberg foundation and operated by NAISS.

## Abbreviations

AI: Artificial intelligence
AL: Active learning
AMP: Antimicrobial peptide
APD: Antimicrobial Peptide Database
CV: Cross validation
DMS: Deep mutational scanning
ESM: Evolutionary scale modeling
GPR: Gaussian process regressor
MIC: Minimal inhibitory concentration
ML: Machine learning
MSA: Multiple sequence alignment
MSE: Mean squared error
PLM: Protein language model
SCARSE: Small-sample Cross-validation-Anchored Regression Solution for low resource peptide Engineering
Std: Standard deviation
TF-IDF: Term frequency–inverse document frequency

## References

1 J. Devlin, M.-W. Chang, K. Lee and K. Toutanova, in Proceedings of the 2019 Conference of the North American Chapter of the Association for Computational Linguistics: Human Language Technologies, Volume 1 (Long and Short Papers), eds J. Burstein, C. Doran and T. Solorio, Association for Computational Linguistics, Minneapolis, Minnesota, 2019, pp. 4171–4186, DOI: 10.18653/v1/N19-1423.

2 A. Radford, K. Narasimhan, T. Salimans and I. Sutskever, Improving language understanding by generative pre-training, OpenAI, 2018.

3 F. Wan, D. Kontogiorgos-Heintz and C. de la Fuente-Nunez, Digit. Discov., 2022, 1, 195–208, DOI: 10.1039/D1DD00024A.

4 R. Schmirler, M. Heinzinger and B. Rost, Nat. Commun., 2024, 15, 7407, DOI: 10.1038/s41467-024-51844-2.

5 S. Biswas, G. Khimulya, E. C. Alley, K. M. Esvelt and G. M. Church, Nat. Methods, 2021, 18, 389–396, DOI: 10.1038/s41592-021-01100-y.

6 E. C. Alley, G. Khimulya, S. Biswas, M. AlQuraishi and G. M. Church, Nat. Methods, 2019, 16, 1315–1322, DOI: 10.1038/s41592-019-0598-1.

7 S. C. Potter, A. Luciani, S. R. Eddy, Y. Park, R. Lopez and R. D. Finn, Nucleic Acids Res., 2018, 46, W200–W204, DOI: 10.1093/nar/gky448.

8 Z. Zhou, L. Zhang, Y. Yu, B. Wu, M. Li, L. Hong and P. Tan, Nat. Commun., 2024, 15, 5566, DOI: 10.1038/s41467-024-49798-6.

9 J.-Y. Chen, J.-F. Wang, Y. Hu, X.-H. Li, Y.-R. Qian and C.-L. Song, Front. Bioeng. Biotechnol., 2025, 13, 1506508, DOI: 10.3389/fbioe.2025.1506508.

10 L. Wang, X. Li, H. Zhang, J. Wang, D. Jiang, Z. Xue and Y. Wang, arXiv, 2025, preprint, DOI: 10.48550/arXiv.2502.06881.

11 P. Notin, A. Kollasch, D. Ritter, L. van Niekerk, S. Paul, H. Spinner, N. Rollins, A. Shaw, R. Orenbuch, R. Weitzman, J. Frazer, M. Dias, D. Franceschi, Y. Gal and D. Marks, in Advances in Neural Information Processing Systems, eds A. Oh, T. Naumann, A. Globerson, K. Saenko, M. Hardt and S. Levine, Curran Associates, Inc., 2023, vol. 36, pp. 64331–64379.

12 M. Li, Y. Tan, X. Ma, B. Zhong, H. Yu, Z. Zhou, W. Ouyang, B. Zhou, P. Tan and L. Hong, in The Thirty-eighth Annual Conference on Neural Information Processing Systems, 2024.

13 J. Su, C. Han, Y. Zhou, J. Shan, X. Zhou and F. Yuan, bioRxiv, 2024, preprint, DOI: 10.1101/2023.10.01.560349.

14 J. Jumper, R. Evans, A. Pritzel, T. Green, M. Figurnov, O. Ronneberger, K. Tunyasuvunakool, R. Bates, A. Žídek, A. Potapenko, A. Bridgland, C. Meyer, S. A. A. Kohl, A. J. Ballard, A. Cowie, B. Romera-Paredes, S. Nikolov, R. Jain, J. Adler, T. Back, S. Petersen, D. Reiman, E. Clancy, M. Zielinski, M. Steinegger, M. Pacholska, T. Berghammer, S. Bodenstein, D. Silver, O. Vinyals, A. W. Senior, K. Kavukcuoglu, P. Kohli and D. Hassabis, Nature, 2021, 596, 583–589, DOI: 10.1038/s41586-021-03819-2.

15 Z. Lin, H. Akin, R. Rao, B. Hie, Z. Zhu, W. Lu, N. Smetanin, R. Verkuil, O. Kabeli, Y. Shmueli, A. dos Santos Costa, M. Fazel-Zarandi, T. Sercu, S. Candido and A. Rives, Science, 2023, 379, 1123–1130, DOI: 10.1126/science.ade2574.

16 C. Guntuboina, A. Das, P. Mollaei, S. Kim and A. Barati Farimani, J. Phys. Chem. Lett., 2023, 14, 10427–10434, DOI: 10.1021/acs.jpclett.3c02398.

17 A. Elnaggar, M. Heinzinger, C. Dallago, G. Rehawi, Y. Wang, L. Jones, T. Gibbs, T. Feher, C. Angerer, M. Steinegger, D. Bhowmik and B. Rost, IEEE Trans. Pattern Anal. Mach. Intell., 2022, 44, 7112–7127, DOI: 10.1109/TPAMI.2021.3095381.

18 S. Badrinarayanan, C. Guntuboina, P. Mollaei and A. Barati Farimani, J. Chem. Inf. Model., 2025, 65, 83–91, DOI: 10.1021/acs.jcim.4c01443.

19 D. Garzon Otero, O. Akbari and C. Bilodeau, Mol. Syst. Des. Eng., 2025, 10, 205–218, DOI: 10.1039/D4ME00172A.

20 D. Garzon Otero, O. Akbari, A. Mandapati and C. Bilodeau, J. Chem. Inf. Model., 2026, 66, 1264–1273, DOI: 10.1021/acs.jcim.5c02629.

21 D. Weininger, J. Chem. Inf. Comput. Sci., 1988, 28, 31–36, DOI: 10.1021/ci00057a005.

22 D. Weininger, A. Weininger and J. L. Weininger, J. Chem. Inf. Comput. Sci., 1989, 29, 97–101, DOI: 10.1021/ci00062a008.

23 M. A. Siani, D. Weininger and J. M. Blaney, J. Chem. Inf. Comput. Sci., 1994, 34, 588–593, DOI: 10.1021/ci00019a017.

24 C.-R. Chung, C.-Y. Chien, Y. Tang, L.-C. Wu, J. B.-K. Hsu, J.-J. Lu, T.-Y. Lee, C. Bai and J.-T. Horng, iScience, 2024, 27, 110718, DOI: 10.1016/j.isci.2024.110718.

25 A. S. Rathore, N. Kumar, S. Choudhury, N. K. Mehta and G. P. S. Raghava, Commun. Biol., 2025, 8, 176, DOI: 10.1038/s42003-025-07615-w.

26 V. I. Levenshtein, Sov. Phys. Dokl., 1965, 10, 707–710.

27 A. T. Müller, G. Gabernet, J. A. Hiss and G. Schneider, Bioinformatics, 2017, 33, 2753–2755, DOI: 10.1093/bioinformatics/btx285.

28 R. Gautier, D. Douguet, B. Antonny and G. Drin, Bioinformatics, 2008, 24, 2101–2102, DOI: 10.1093/bioinformatics/btn392.

29 R. C. A. Keller, Int. J. Mol. Sci., 2011, 12, 5577–5591, DOI: 10.3390/ijms12095577.

30 F. Pedregosa, G. Varoquaux, A. Gramfort, V. Michel, B. Thirion, O. Grisel, M. Blondel, P. Prettenhofer, R. Weiss, V. Dubourg, J. Vanderplas, A. Passos, D. Cournapeau, M. Brucher, M. Perrot and E. Duchesnay, J. Mach. Learn. Res., 2011, 12, 2825–2830.

31 C. E. Rasmussen and C. K. I. Williams, Gaussian processes for machine learning, MIT Press, Cambridge, Mass., 3rd edn., 2008.

32 T. Akiba, S. Sano, T. Yanase, T. Ohta and M. Koyama, in Proceedings of the 25th ACM SIGKDD International Conference on Knowledge Discovery & Data Mining, Association for Computing Machinery, New York, NY, USA, 2019, pp. 2623–2631, DOI: 10.1145/3292500.3330701.

33 J. R. Ash, C. Wognum, R. Rodríguez-Pérez, M. Aldeghi, A. C. Cheng, D.-A. Clevert, O. Engkvist, C. Fang, D. J. Price, J. M. Hughes-Oliver and W. P. Walters, J. Chem. Inf. Model., 2025, 65, 9398–9411, DOI: 10.1021/acs.jcim.5c01609.

34 G. Wang, C. Schmidt, X. Li and Z. Wang, Nucleic Acids Res., 2026, 54, D363–D374, DOI: 10.1093/nar/gkaf860.

