## Supplementary file for "AI-guided discovery for low-resource peptide engineering using evolutionary scale modeling"

### Electronic Supplementary Information (ESI) for: AI-guided discovery for low-resource peptide engineering using evolutionary scale modeling

Leo Andrekson[a,b], Robin Rydbergh[a,b], Rocío Mercado[c,d,*], Michaela Wenzel[a,b,*]

[a]Division of Molecular Bioscience, Department of Life Sciences, Chalmers University of Technology, Gothenburg, Sweden

[b]Centre for Antibiotic Resistance Research in Gothenburg (CARe), Gothenburg, Sweden

[c]Division of Data Science and AI, Department of Computer Science and Engineering, Chalmers University of Technology & University of Gothenburg, Gothenburg, Sweden

[d]Science for Life Laboratory (SciLifeLab), Gothenburg, Sweden

#### Section S1. Datasets

Datasets used for substitutions are summarized below in Table S1.

Table S1. Substitution datasets: the table provides dataset names, source organism, sequence length, total number of mutants, number of one-point mutants, number of two-point mutants, endpoint used for evaluating performance, and range of endpoint values. Stability stands for assays that measure how thermostable a protein or peptide is, while activity stands for assays that directly or indirectly measure a protein/peptide's catalytic (or otherwise biochemical) activity.

| Dataset | Source organism | Sequence length | No. of mutants | No. of one-point mutants | No. of two-point mutants | Endpoint | Range |
|---|---|---|---|---|---|---|---|
| YNZC BACSU sub | *Bacillus subtilis* | 39 aa | 2300 | 714 | 1586 | Stability | [-3.15, 1.38] |
| MAFG MOUSE sub | *Mus musculus* | 41 aa | 1429 | 762 | 667 | Stability | [-2.89, 0.92] |
| SDA BACSU | *Bacillus subtilis* | 44 aa | 2770 | 834 | 1936 | Stability | [-4.41, 1.20] |
| POLG PESV sub | Porcine enteric sapovirus | 53 aa | 5130 | 995 | 4135 | Stability | [-4.90, 1.04] |
| DN7A SACS2 | *Saccharolobus solfataricus* | 55 aa | 1008 | 1008 | - | Stability | [-5.16, 0.69] |
| SBI STAAM | *Staphylococcus aureus* | 56 aa | 1025 | 1025 | - | Stability | [-2.30, 1.01] |
| SOX30 HUMAN | *Homo sapiens* | 57 aa | 1010 | 1010 | - | Stability | [-3.81, 1.71] |

| | | | | | | | |
|---|---|---|---|---|---|---|---|
| ENVZ ECOLI | *Escherichia coli* | 60 aa | 1121 | 1121 | - | Activity | [0.80, 4.70] |
| FKBP3 HUMAN | *Homo sapiens* | 69 aa | 1237 | 1237 | - | Stability | [-3.29, 1.90] |
| A0A247D711 LISMN | *Listeria monocytogenes* | 87 aa | 1653 | 1653 | - | Activity | [-4.84, 0.96] |

In addition, we used the following datasets for indels, summarized in Table S2.

Table S2. Indel datasets: the table provides dataset name, source organism, WT sequence length, number of mutants, mean Levenshtein distance from wild type (WT) across mutants, endpoint used for evaluating performance, and range of endpoint values [20]. Stability stands for assays that measure how thermostable a protein or peptide is.

| Dataset | Source organism | WT sequence length | No. of mutants | Mean Levenshtein distance from WT | Endpoint | Range |
|---|---|---|---|---|---|---|
| PIN1 HUMAN | *Homo sapiens* | 39 aa | 106 | 1 | Stability | [-2.86, 1.18] |
| YNZC BACSU ind | *Bacillus subtilis* | 39 aa | 104 | 1 | Stability | [-3.12, 0.28] |
| VG08 BPP22 | *Salmonella* phage P22 | 40 aa | 101 | 1 | Stability | [-2.02, 0.35] |
| SQSTM MOUSE | *Mus musculus* | 40 aa | 111 | 1 | Stability | [-2.49, 0.30] |
| MAFG MOUSE ind | *Mus musculus* | 41 aa | 115 | 1 | Stability | [-2.62, 0.38] |
| RD23A HUMAN | *Homo sapiens* | 44 aa | 120 | 1 | Stability | [-3.79, 0.00] |
| SDA BACSU ind | *Bacillus subtilis* | 44 aa | 127 | 1 | Stability | [-4.20, 0.12] |
| POLG PESV ind | Porcine enteric sapovirus | 53 aa | 149 | 1 | Stability | [-4.77, 0.42] |
| RPC1 BP434 | Enterobacteria phage 434 | 61 aa | 164 | 1 | Stability | [-4.04, 1.14] |
| PKN1 HUMAN | *Homo sapiens* | 71 aa | 187 | 1 | Stability | [-3.26, 0.48] |

The final four datasets consist of the *Escherichia coli*, *Staphylococcus aureus*, and *Pseudomonas aeruginosa* AMPs datasets and the HemoPI2 dataset (see Table S3).

Table S3. Additional datasets: the table provides dataset name, sequence length range, number of sequences, endpoint used for evaluating performance, and range of endpoint values. Endpoint of the *E. coli*, *S. aureus*, and *P. aeruginosa* AMPs datasets, pMIC, corresponds to the negative logarithm of MIC, while the $pHC_{50}$ of HemoPI2 corresponds to the negative logarithm of $HC_{50}$.

| Dataset | Sequence length range | No. of sequences | Endpoint | Range |
|---|---|---|---|---|
| *E. coli* AMPs | 6-40 aa | 3818 | pMIC | [-3.67, 2.30] |
| *S. aureus* AMPs | 6-40 aa | 2644 | pMIC | [-4.01, 3.33] |
| *P. aeruginosa* AMPs | 6-40 aa | 2458 | pMIC | [-3.03, 2.40] |
| HemoPI2 | 6-39 aa | 1926 | $pHC_{50}$ | [-3.00,0.72] |

Endpoint distribution of all substitution datasets in Table S1 are visualized in Fig. S1.

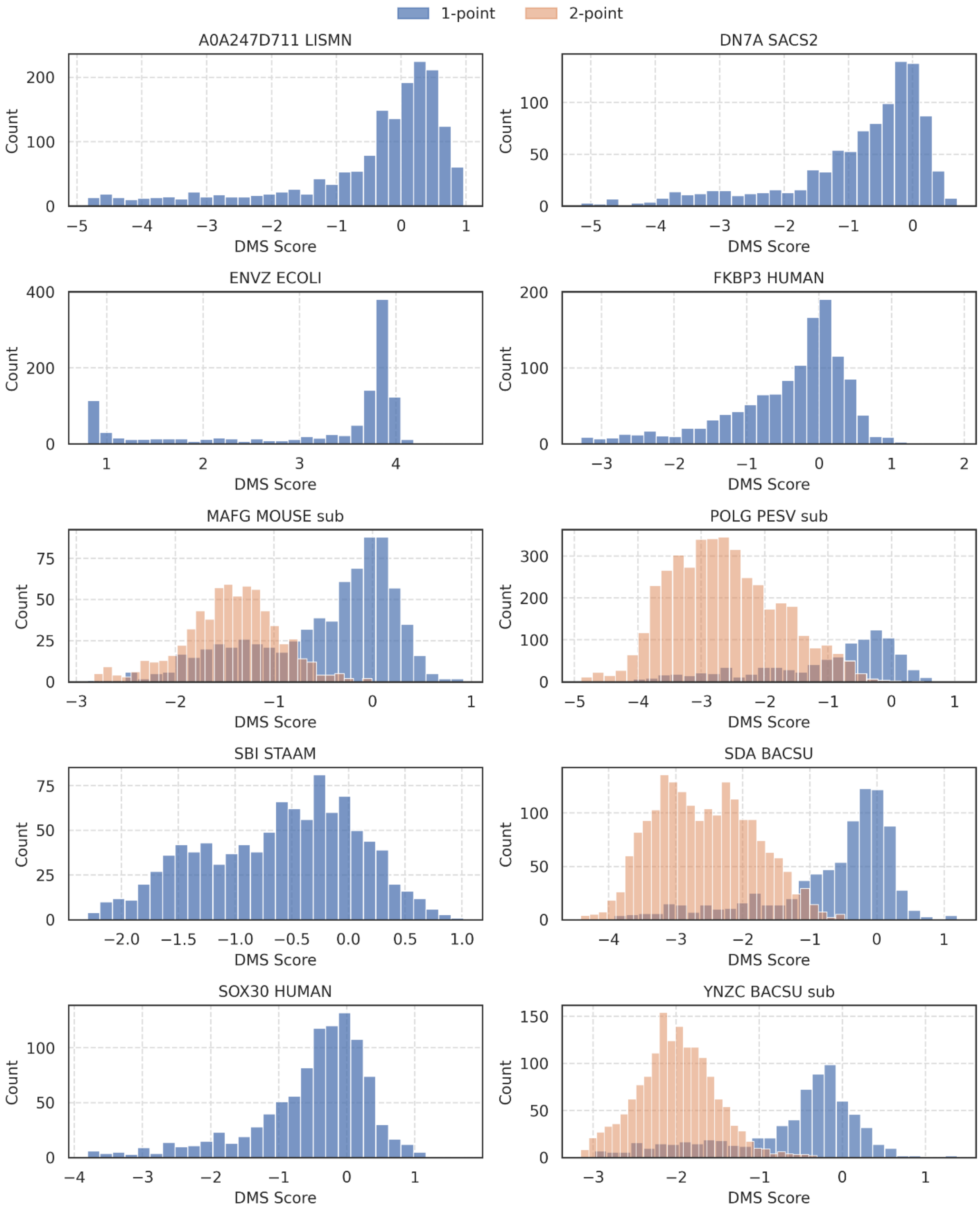


Fig. S1. Substitution-based datasets. Distribution of DMS score for all 10 substitution-based datasets in Table S1, colored by one and two-point mutants.

Similarly, all indel based datasets in Table S2 are visualized in Fig. S2.

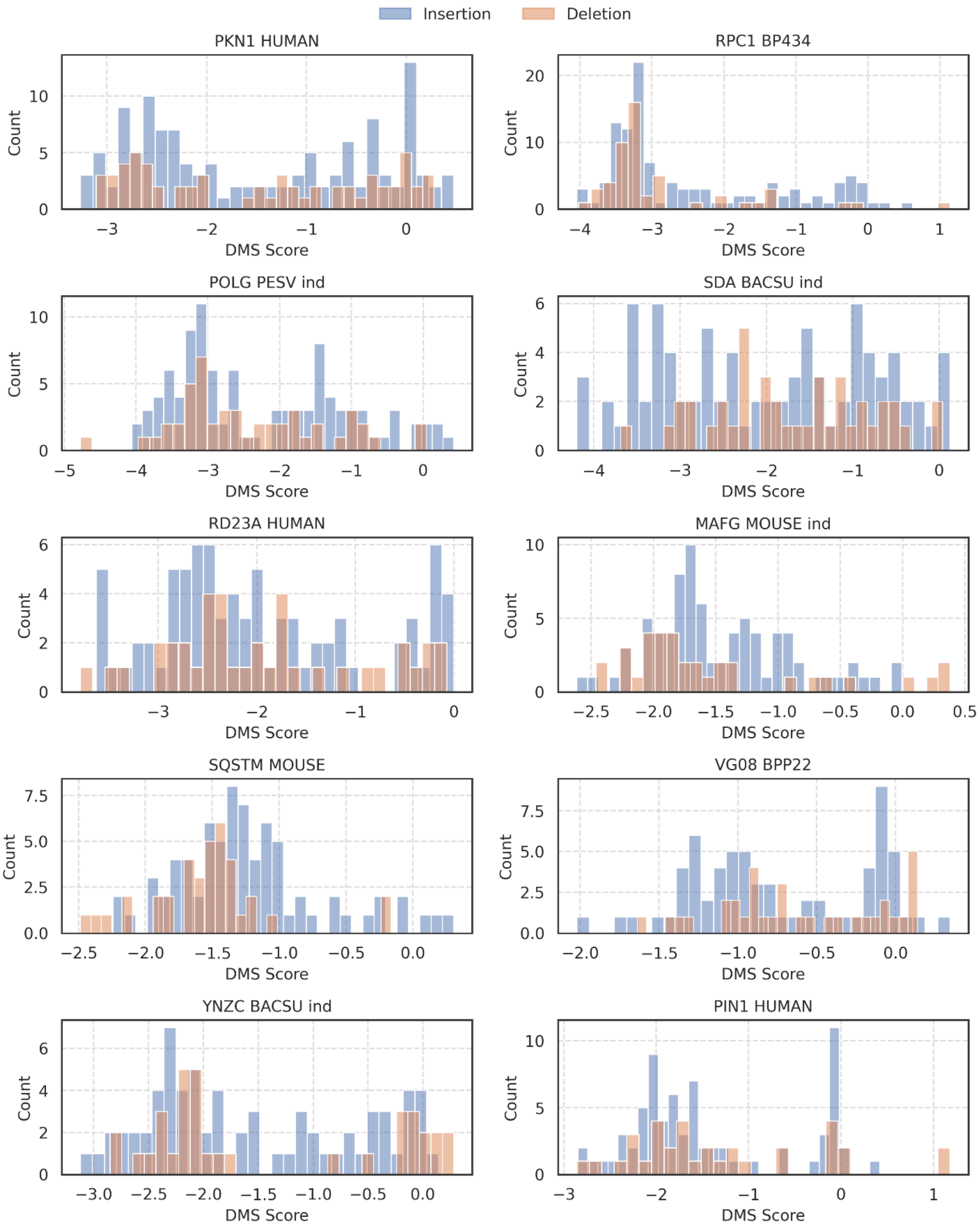


Fig. S2. Indel-based datasets. Distribution of DMS score for all 10 indel datasets in Table S2, colored by insertion and deletion-based mutants.

The final four datasets from Table S3 are visualized below (Fig. S3).

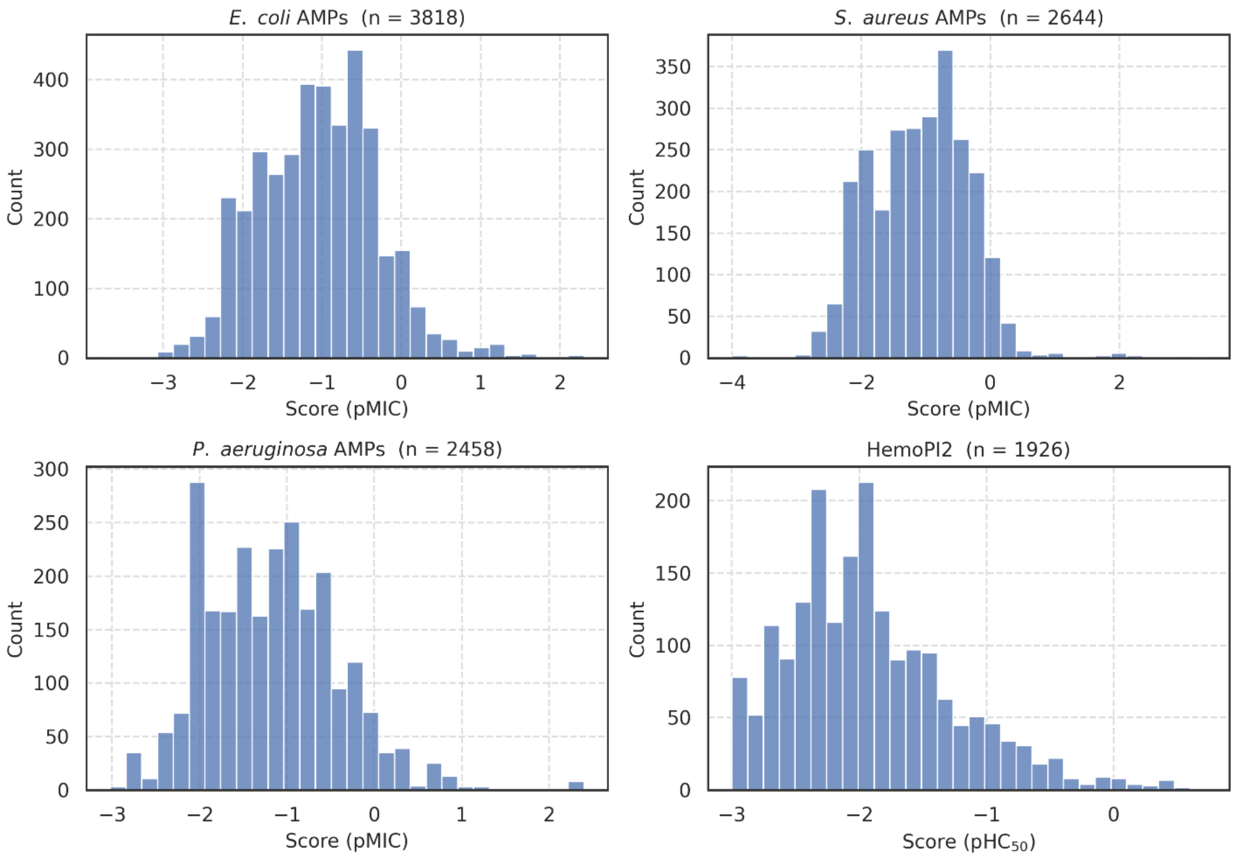


Fig. S3. *E. coli* AMPs, *S. aureus* AMPs, *P. aeruginosa* AMPs, and the HemoPI2 dataset. Distribution of pMIC score for the sequences in the AMP datasets, together with the distribution of $pHC_{50}$ score for the HemoPI2 dataset.

#### Section S2. ESM2 embedding evaluation

An initial pre-screen was performed to assess which ESM2 foundation model to use for SCARSE. The 8, 35, 150, and 650 million parameter ESM2 models were evaluated for the substitution and indel datasets, in addition to the remaining 4 peptide datasets under the group name "Peptides" in Fig. S4.

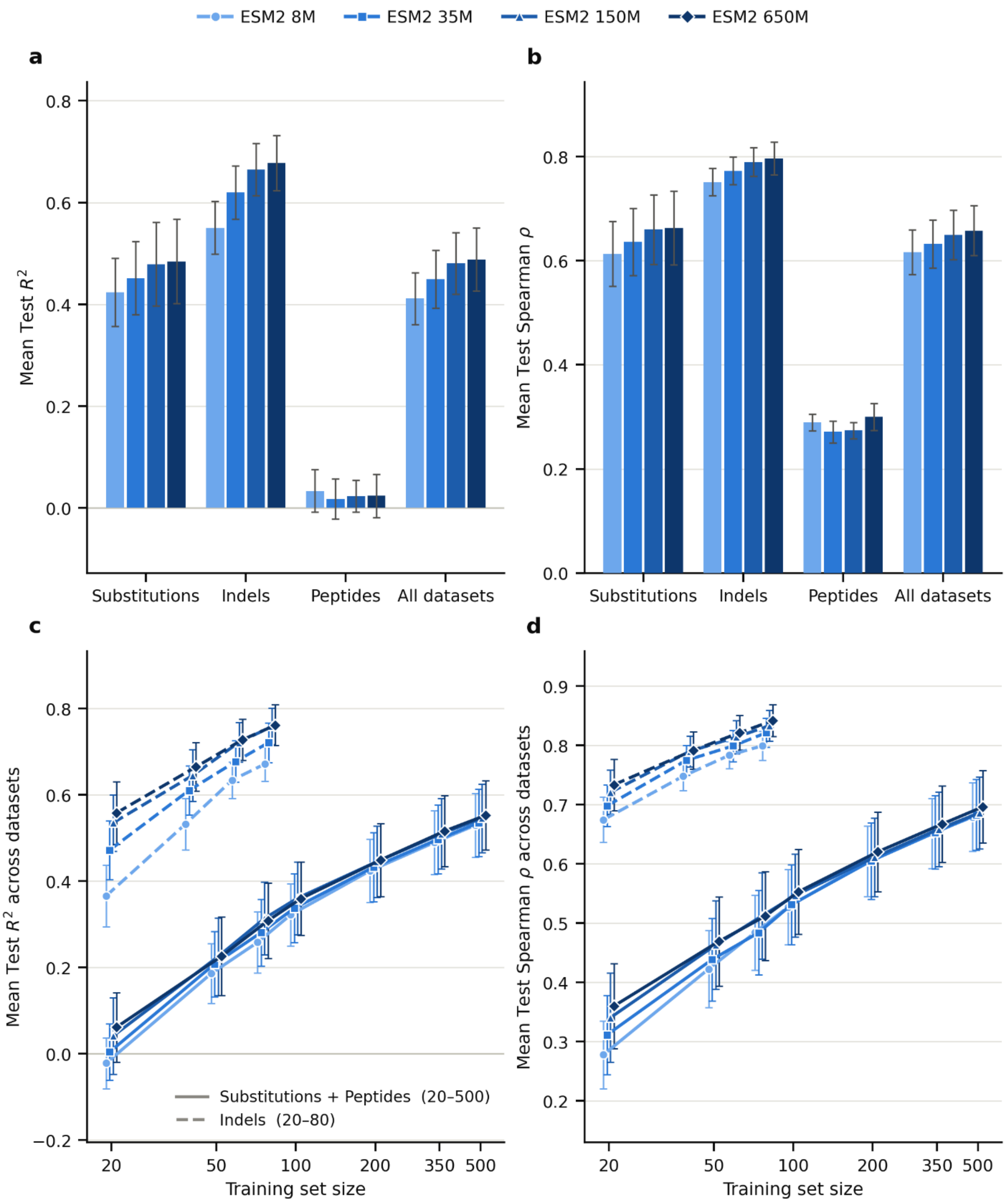


Fig. S4. Effect of ESM2 model size on predictive performance. Gaussian process regression (GPR) models were trained on embeddings from four ESM2 checkpoints (8M, 35M, 150M and 650M parameters). They were evaluated on the substitution (n = 10), indel (n = 10) and peptide (n = 4) datasets, with 10 random seeds per training-set size per dataset. The peptide group comprises three antimicrobial-peptide datasets, with potency against *E. coli*, *S. aureus* and *P. aeruginosa* measured as pMIC, and the HemoPI2 hemolysis dataset ($pHC_{50}$). (a, b) Mean test $R^2$ (a) and mean test Spearman ρ (b) per checkpoint for each dataset family and across all 24 datasets. Each dataset contributes one value, averaged over seeds and training-set sizes, so families with more training-set sizes are not weighted more heavily. Error bars show ±1 standard error of the mean (s.e.m.) across datasets. (c, d) Mean test $R^2$ (c) and mean test Spearman ρ (d) across datasets as a function of training-set size (log scale). Solid lines average the substitution and peptide datasets, which share a 20–500 training-size grid. Dashed lines average the indel datasets, evaluated at 20–80. Error bars show ±1 s.e.m. across datasets. Markers are offset slightly horizontally for legibility.

Performance increased with model size for both the substitution and indel datasets (Fig. S4a, b). Averaged over all datasets, mean test $R^2$ rose from approximately 0.41 for the 8M model to 0.45, 0.48 and 0.49 for the 35M, 150M and 650M models, respectively, with the same ordering for Spearman ρ (0.62 to 0.66). The gain was largest for the indel datasets, where mean $R^2$ increased from approximately 0.55 (8M) to 0.68 (650M), compared with 0.42 to 0.49 for the substitution datasets. For the peptide datasets, all four models performed similarly, with mean $R^2$ close to zero and Spearman ρ between approximately 0.27 and 0.30, showing no consistent effect of model size.

The benefit of a larger model depended on the amount of training data (Fig. S4c, d). For the indel datasets, the difference between the smallest and largest model was greatest at the smallest training-set size, approximately 0.37 versus 0.56 in $R^2$ at 20 training examples, and narrowed as more data became available. Similarly, for the substitution and peptide datasets, the larger models held a small advantage at 20–75 training examples, whereas the four models converged from approximately 200 training examples onwards. Differences between the 150M and 650M models were small throughout. The error bars overlapped for adjacent model sizes, indicating that the variation between datasets exceeded the differences between models.

Because the 650M model performed best, or equal to best, in every dataset family and for both metrics, it was selected as the foundation model for SCARSE and used in all subsequent analyses.

#### Section S3. Model hyperparameters

Below, we provide hyperparameters used for optimization of the gaussian process regressor (GPR) (Table S4).

Table S4. Hyperparameter optimization space: the table contains top model name, parameter name, which type, i.e., continuous (Cont) or categorical (Cat), and the options for selection or range of continuous values.

| Model | Parameter | Type | Range/Options |
|---|---|---|---|
| GPR | Alpha | Cont | [$1e^{-10}$, $1e^{-6}$] |
| | Normalize y | Cat | [True, False] |
| | Kernel | Cat | [rbf, matern, rationalquadratic, dotproduct] |

The three baseline representations used for comparison against SCARSE (the Methods (Input representation)) are summarized in Table S5.

Table S5. Baseline representations: the table contains descriptions of what defines each baseline. Each baseline is combined with the same Gaussian process regressor (GPR) as SCARSE.

| Representation | Description | Settings |
|---|---|---|

| | | |
|---|---|---|
| Physchem | modlAMP global descriptors together with amino acid composition, the fractions of polar, apolar, charged and aromatic residues, the hydrophobic moment, and the discrimination factor. | As described in the Methods (Input representation). |
| Morgan | Morgan (circular) fingerprint computed from the peptide's molecular structure. | radius = 2, length = 2048 bits (binary circular fingerprint; molecule built from the sequence with RDKit MolFromSequence). |
| N-gram | Normalized counts of contiguous amino acid subsequences (character n-grams) of the sequence. | Character n-grams, order 1–3 (ngram_range = (1, 3)), TF-IDF weighted with a vocabulary fit on the training sequences only. |

#### Section S4. Benchmarking performance

Fig. S5a–j reports the performance of SCARSE (GPR + ESM2) next to the three baseline descriptors (Physchem, Morgan, and N-gram) for all substitution datasets. We display the mean $R^2$ value of the test sets across 10 random seeds per dataset and training size together with the standard deviation (std) error bars. Note that a lower limit of -10 was set for the test $R^2$ of the SDA BACSU dataset for visualization due to instabilities occurring for the Physchem baseline at training sizes 50, 75, and 100. A total of 5 seeds were filtered out by this, with the lowest seed reaching a $R^2$ of -312. This was done to improve the visualizations, but the statistics were still carried out without this filter to ensure a representative statistical analysis.

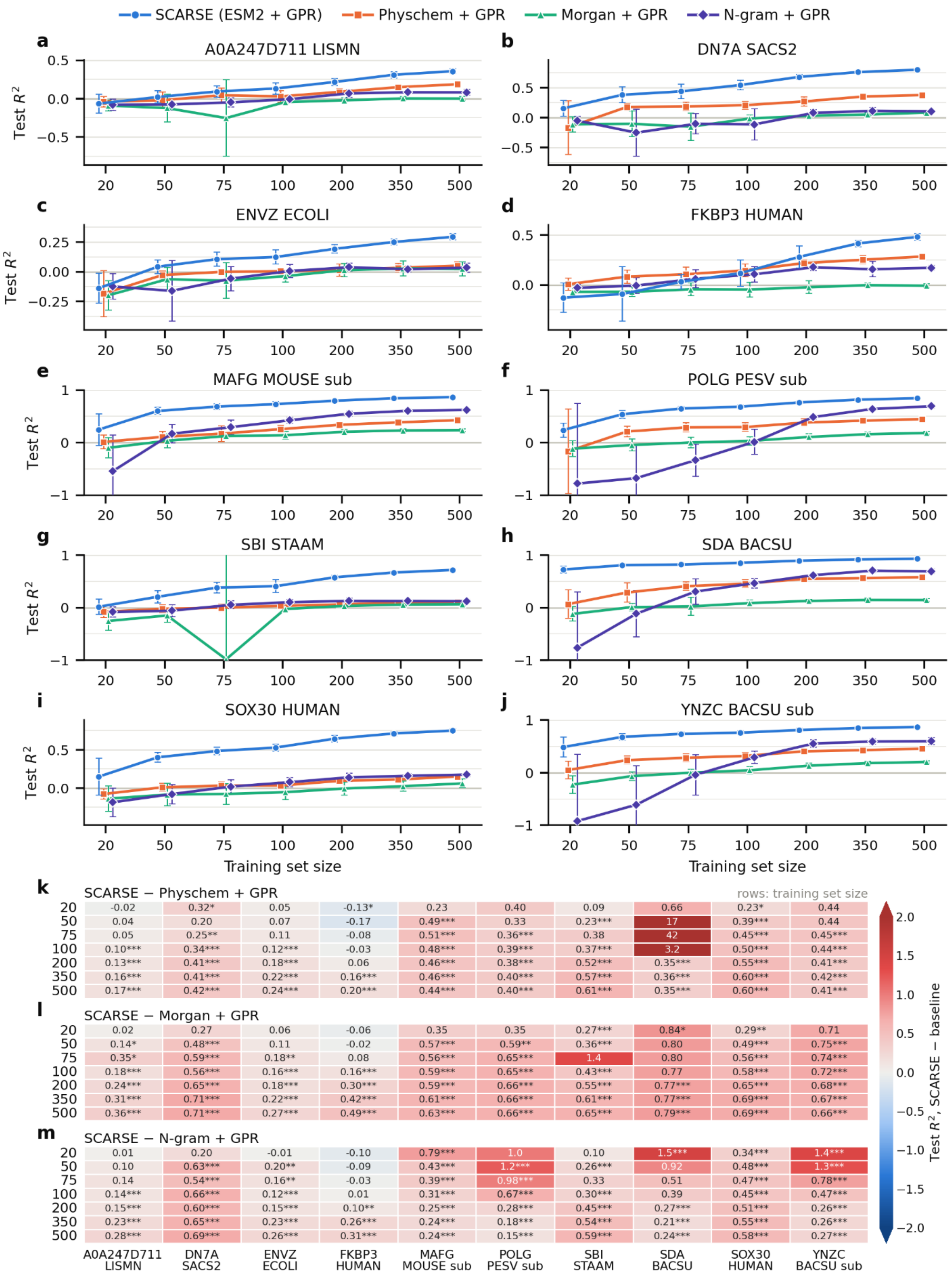
SCARSE (ESM2 + GPR)
Physchem + GPR
Morgan + GPR
N-gram + GPR
a
A0A247D711 LISMN
b
DN7A SACS2
c
ENVZ ECOLI
d
FKBP3 HUMAN
e
MAFG MOUSE sub
f
POLG PESV sub
g
SBI STAAM
h
SDA BACSU
i
SOX30 HUMAN
j
YNZC BACSU sub
Test $R^2$
Training set size
20 50 75 100 200 350 500
k
SCARSE − Physchem + GPR
rows: training set size
20: -0.02 0.32* 0.05 -0.13* 0.23 0.40 0.09 0.66 0.23* 0.44
50: 0.04 0.20 0.07 -0.17 0.49*** 0.33 0.23*** 17 0.39*** 0.44
75: 0.05 0.25** 0.11 -0.08 0.51*** 0.36*** 0.38 42 0.45*** 0.45***
100: 0.10*** 0.34*** 0.12*** -0.03 0.48*** 0.39*** 0.37*** 3.2 0.50*** 0.44***
200: 0.13*** 0.41*** 0.18*** 0.06 0.46*** 0.38*** 0.52*** 0.35*** 0.55*** 0.41***
350: 0.16*** 0.41*** 0.22*** 0.16*** 0.46*** 0.40*** 0.57*** 0.36*** 0.60*** 0.42***
500: 0.17*** 0.42*** 0.24*** 0.20*** 0.44*** 0.40*** 0.61*** 0.35*** 0.60*** 0.41***
l
SCARSE − Morgan + GPR
20: 0.02 0.27 0.06 -0.06 0.35 0.35 0.27*** 0.84* 0.29** 0.71
50: 0.14* 0.48*** 0.11 -0.02 0.57*** 0.59** 0.36*** 0.80 0.49*** 0.75***
75: 0.35* 0.59*** 0.18** 0.08 0.56*** 0.65*** 1.4 0.80 0.56*** 0.74***
100: 0.18*** 0.56*** 0.16*** 0.16*** 0.59*** 0.65*** 0.43*** 0.77 0.58*** 0.72***
200: 0.24*** 0.65*** 0.18*** 0.30*** 0.59*** 0.66*** 0.55*** 0.77*** 0.65*** 0.68***
350: 0.31*** 0.71*** 0.22*** 0.42*** 0.61*** 0.66*** 0.61*** 0.77*** 0.69*** 0.67***
500: 0.36*** 0.71*** 0.27*** 0.49*** 0.63*** 0.66*** 0.65*** 0.79*** 0.69*** 0.66***
m
SCARSE − N-gram + GPR
20: 0.01 0.20 -0.01 -0.10 0.79*** 1.0 0.10 1.5*** 0.34*** 1.4***
50: 0.10 0.63*** 0.20** -0.09 0.43*** 1.2*** 0.26*** 0.92 0.48*** 1.3***
75: 0.14 0.54*** 0.16** -0.03 0.39*** 0.98*** 0.33 0.51 0.47*** 0.78***
100: 0.14*** 0.66*** 0.12*** 0.01 0.31*** 0.67*** 0.30*** 0.39 0.45*** 0.47***
200: 0.15*** 0.60*** 0.15*** 0.10** 0.25*** 0.28*** 0.45*** 0.27*** 0.51*** 0.26***
350: 0.23*** 0.65*** 0.23*** 0.26*** 0.24*** 0.18*** 0.54*** 0.21*** 0.55*** 0.26***
500: 0.28*** 0.69*** 0.26*** 0.31*** 0.24*** 0.15*** 0.59*** 0.24*** 0.58*** 0.27***
A0A247D711 LISMN
DN7A SACS2
ENVZ ECOLI
FKBP3 HUMAN
MAFG MOUSE sub
POLG PESV sub
SBI STAAM
SDA BACSU
SOX30 HUMAN
YNZC BACSU sub
Test $R^2$, SCARSE − baseline
2.0 1.5 1.0 0.5 0.0 −0.5 −1.0 −1.5 −2.0

Fig. S5. Benchmarking performance of substitution datasets (Test $R^2$). (a–j) Benchmarking performance in terms of mean $R^2 \pm$ std across 10 random seeds. (k–m) Effect-size difference between the means of SCARSE and each baseline together with the Tukey HSD post-hoc statistical test (*$p \leq 0.05$, **$p \leq 0.01$, ***$p \leq 0.001$).

In general, the performance of the models increases with the number of samples used for training. Notably, for many datasets (Fig. S5b, e–j), relatively high $R^2$ values can be achieved with only a small number of data points, suggesting that applying SCARSE may be beneficial even in low-sample settings. Fig. S5k–m illustrates the effect-size difference between the means of SCARSE and the baselines together with the statistical significance. For most datasets and training sizes, SCARSE is statistically significantly better than the baselines, highlighting the improvement potential from using ESM2. The corresponding Spearman ρ rank correlation is shown in Fig. S6.

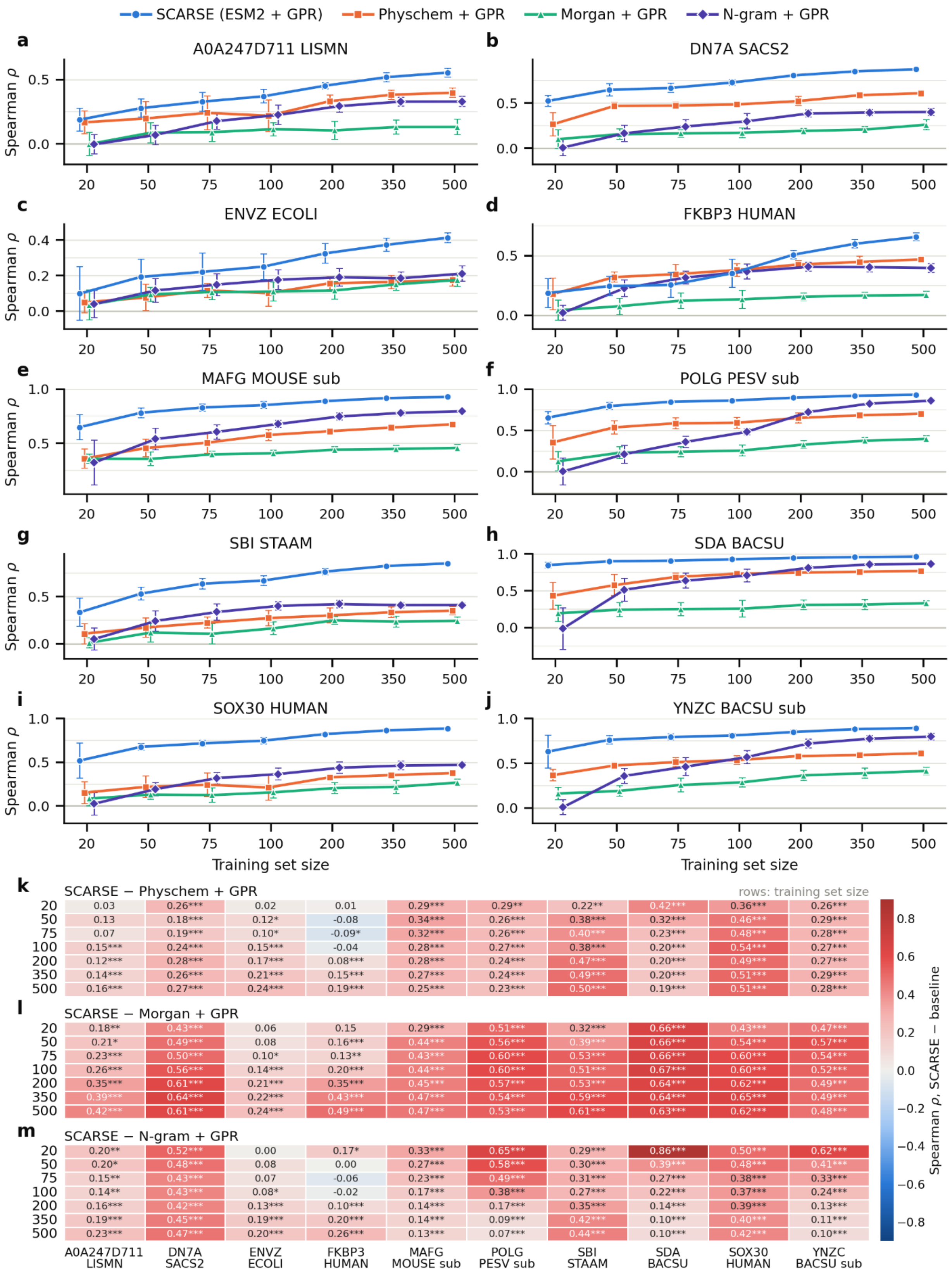

SCARSE (ESM2 + GPR)
Physchem + GPR
Morgan + GPR
N-gram + GPR
a
A0A247D711 LISMN
b
DN7A SACS2
c
ENVZ ECOLI
d
FKBP3 HUMAN
e
MAFG MOUSE sub
f
POLG PESV sub
g
SBI STAAM
h
SDA BACSU
i
SOX30 HUMAN
j
YNZC BACSU sub
Spearman ρ
Training set size
20 50 75 100 200 350 500
k
SCARSE − Physchem + GPR
rows: training set size
20 0.03 0.26*** 0.02 0.01 0.29*** 0.29** 0.22** 0.42*** 0.36*** 0.26***
50 0.13 0.18*** 0.12* -0.08 0.34*** 0.26*** 0.38*** 0.32*** 0.46*** 0.29***
75 0.07 0.19*** 0.10* -0.09* 0.32*** 0.26*** 0.40*** 0.23*** 0.48*** 0.28***
100 0.15*** 0.24*** 0.15*** -0.04 0.28*** 0.27*** 0.38*** 0.20*** 0.54*** 0.27***
200 0.12*** 0.28*** 0.17*** 0.08*** 0.28*** 0.24*** 0.47*** 0.20*** 0.49*** 0.27***
350 0.14*** 0.26*** 0.21*** 0.15*** 0.27*** 0.24*** 0.49*** 0.20*** 0.51*** 0.29***
500 0.16*** 0.27*** 0.24*** 0.19*** 0.25*** 0.23*** 0.50*** 0.19*** 0.51*** 0.28***
l
SCARSE − Morgan + GPR
20 0.18** 0.43*** 0.06 0.15 0.29*** 0.51*** 0.32*** 0.66*** 0.43*** 0.47***
50 0.21* 0.49*** 0.08 0.16*** 0.44*** 0.56*** 0.39*** 0.66*** 0.54*** 0.57***
75 0.23*** 0.50*** 0.10* 0.13** 0.43*** 0.60*** 0.53*** 0.66*** 0.60*** 0.54***
100 0.26*** 0.56*** 0.14*** 0.20*** 0.44*** 0.60*** 0.51*** 0.67*** 0.60*** 0.52***
200 0.35*** 0.61*** 0.21*** 0.35*** 0.45*** 0.57*** 0.53*** 0.64*** 0.62*** 0.49***
350 0.39*** 0.64*** 0.22*** 0.43*** 0.47*** 0.54*** 0.59*** 0.64*** 0.65*** 0.49***
500 0.42*** 0.61*** 0.24*** 0.49*** 0.47*** 0.53*** 0.61*** 0.63*** 0.62*** 0.48***
m
SCARSE − N-gram + GPR
20 0.20** 0.52*** 0.00 0.17* 0.33*** 0.65*** 0.29*** 0.86*** 0.50*** 0.62***
50 0.20* 0.48*** 0.08 0.00 0.27*** 0.58*** 0.30*** 0.39*** 0.48*** 0.41***
75 0.15** 0.43*** 0.07 -0.06 0.23*** 0.49*** 0.31*** 0.27*** 0.38*** 0.33***
100 0.14** 0.43*** 0.08* -0.02 0.17*** 0.38*** 0.27*** 0.22*** 0.37*** 0.24***
200 0.16*** 0.42*** 0.13*** 0.10*** 0.14*** 0.17*** 0.35*** 0.14*** 0.39*** 0.13***
350 0.19*** 0.45*** 0.19*** 0.20*** 0.14*** 0.09*** 0.42*** 0.10*** 0.40*** 0.11***
500 0.23*** 0.47*** 0.20*** 0.26*** 0.13*** 0.07*** 0.44*** 0.10*** 0.42*** 0.10***
A0A247D711 LISMN
DN7A SACS2
ENVZ ECOLI
FKBP3 HUMAN
MAFG MOUSE sub
POLG PESV sub
SBI STAAM
SDA BACSU
SOX30 HUMAN
YNZC BACSU sub
Spearman ρ, SCARSE − baseline

Fig. S6. Benchmarking performance of substitution datasets (Spearman ρ). (a–j) Test Spearman ρ (mean ± std across 10 random seeds) for SCARSE and the three baselines. (k–m) Effect-size difference between the means of SCARSE and each baseline together with the Tukey HSD post-hoc statistical test (*$p \leq 0.05$, **$p \leq 0.01$, ***$p \leq 0.001$).

For the substitution datasets we further illustrate the hypothetical scenario "among the top 20 sequences ranked by predicted performance, the proportion that truly belong to the top 20 based on actual performance" (Fig. S7).

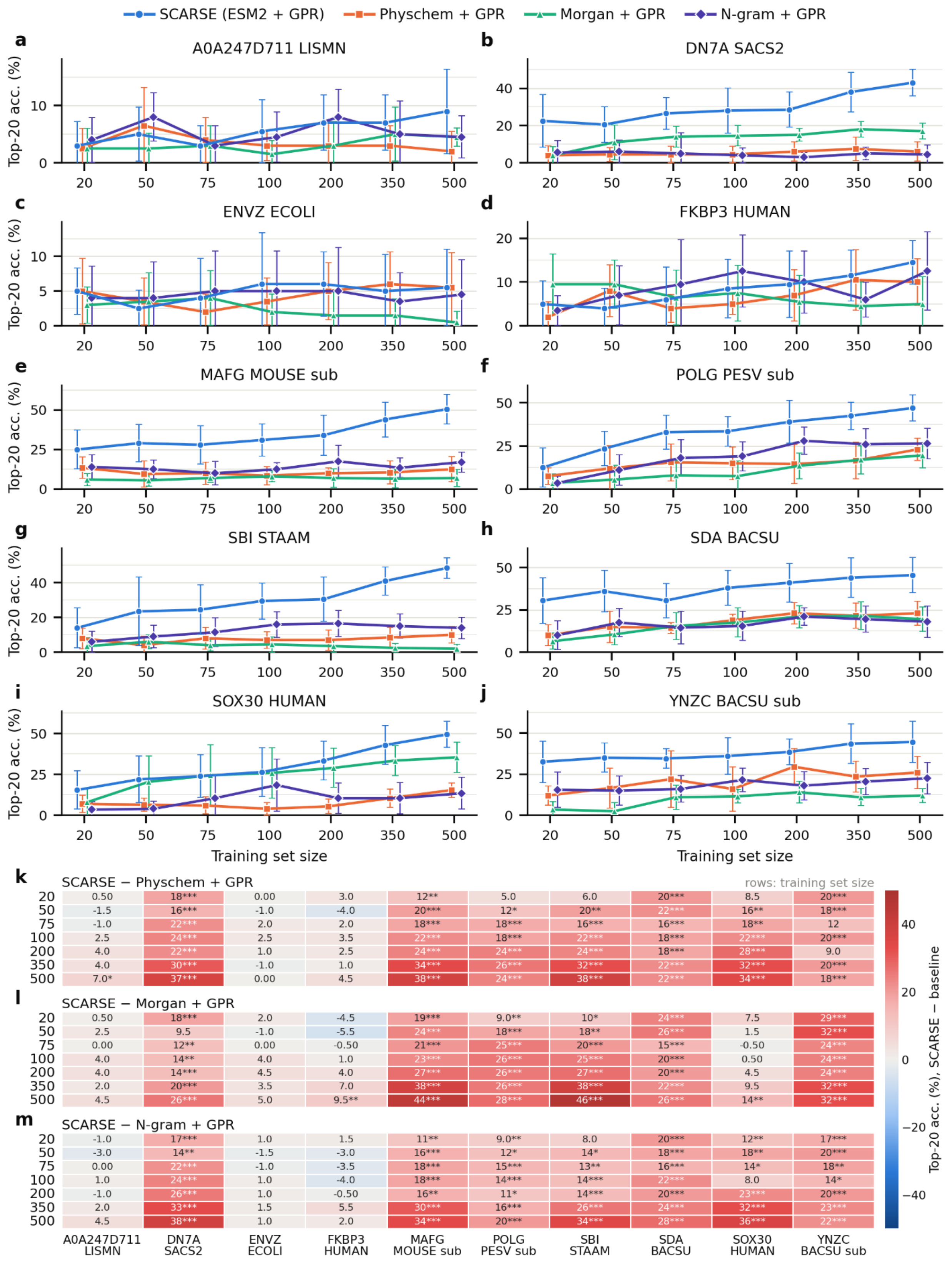


Fig. S7. High-performing selection performance for substitution datasets. (a–j) Benchmarking results evaluated by the following metric: among the top 20 sequences ranked by predicted performance, the proportion that truly belong to the top 20 based on actual performance. Values are reported as mean ± std across 10 random seeds. (k–m) Effect-size difference between the means of SCARSE and each baseline together with the Tukey HSD post-hoc statistical test (*$p \leq 0.05$, **$p \leq 0.01$, ***$p \leq 0.001$).

If one were to randomly select samples, one would expect to observe an average of 4% of these to be in the actual top 20, due to there being a total of 500 peptides in each test set. What we observe in Fig. S7 is that in most cases the percentage is higher than 4% for SCARSE.

In Fig. S8a–j we can see the performance of SCARSE next to the three baselines for all indel datasets.

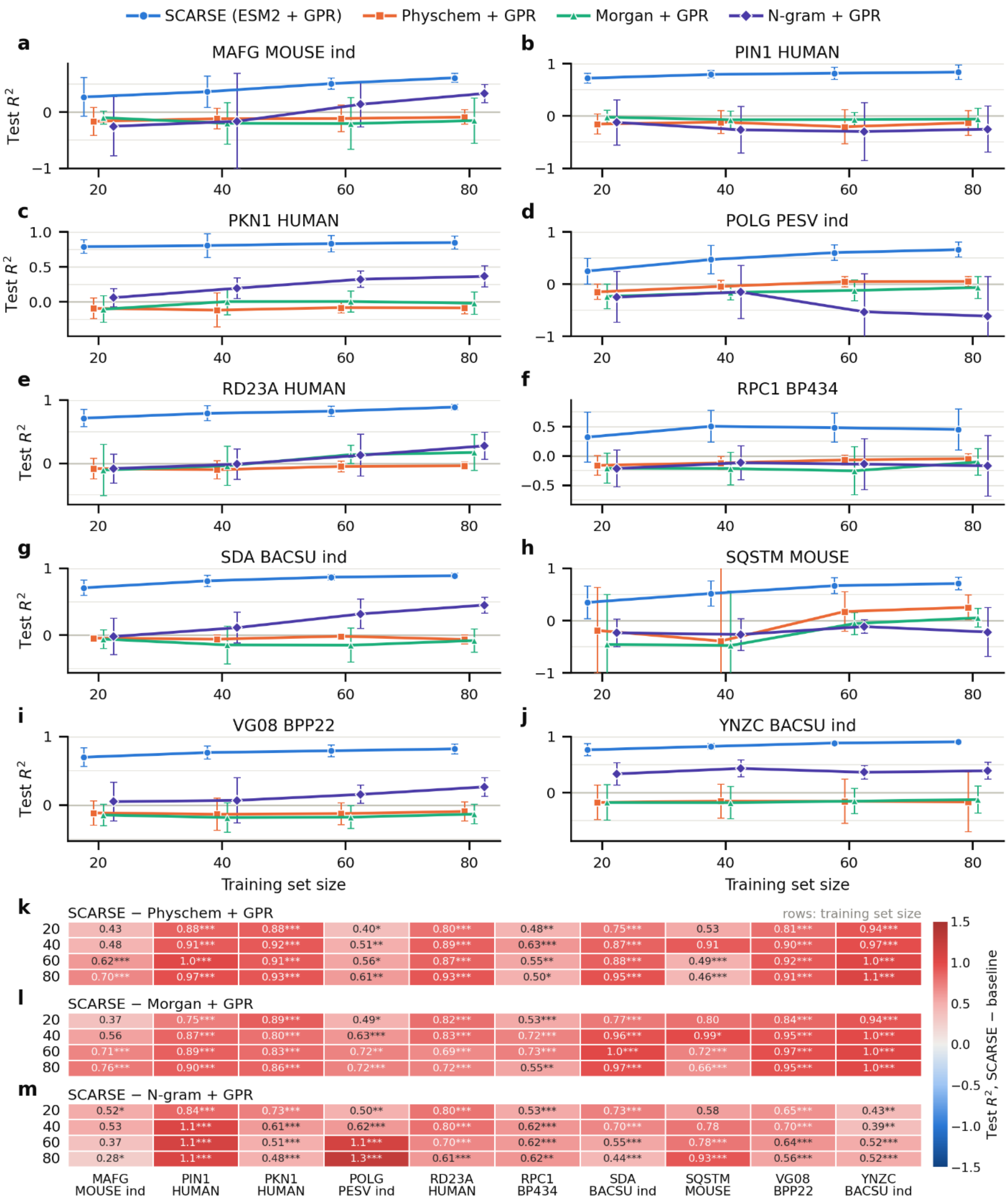


Fig. S8. Benchmarking performance of indel datasets (Test R²). (a–j) Benchmarking performance in terms of mean R² ± std across 10 random seeds. (k–m) Effect-size difference between the means of SCARSE and each baseline together with the Tukey HSD post-hoc statistical test (*p ≤ 0.05, **p ≤ 0.01, ***p ≤ 0.001).

From Fig. S8a–m, we observe a clear improvement from using SCARSE compared to the baseline descriptors. This indicates that ESM2 captures the subtle differences between indel mutants required for generalization, while the baselines often achieve an $R^2$ around 0, showing a lack of information in the baseline descriptors needed for prediction on unseen indels.

Finally, we observe benchmarking results for the *E. coli* AMPs, *S. aureus* AMPs, *P. aeruginosa* AMPs, and the HemoPI2 datasets (Fig. S9a–d).

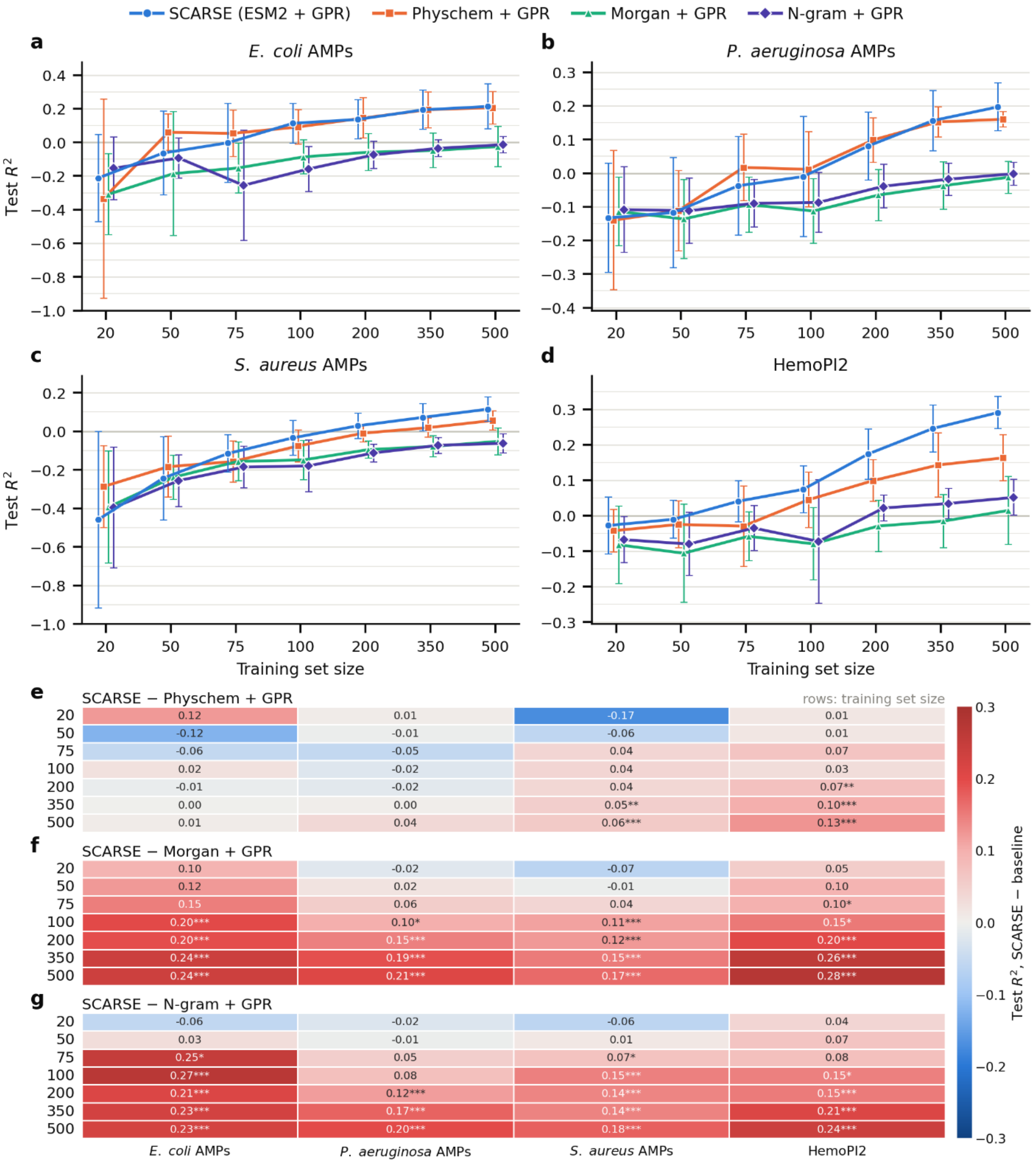


Fig. S9. Benchmarking performance of the peptide datasets (Test $R^2$). (a–d) Benchmarking performance of the *E. coli* AMPs, HemoPI2, *P. aeruginosa* AMPs, and *S. aureus* AMPs datasets in terms of mean $R^2 \pm$ std across 10 random seeds. (e–g) Effect-size difference between the means of SCARSE and each baseline together with the Tukey HSD post-hoc statistical test (*$p \leq 0.05$, **$p \leq 0.01$, ***$p \leq 0.001$).

From Fig. S9e–g we conclude that SCARSE is statistically better than the Morgan and N-gram baselines. When compared to the Physchem baseline, SCARSE has a slight advantage at larger training sizes for the HemoPI2 and S. aureus AMPs datasets, but is generally not statistically better. Looking at the Spearman correlation (Fig. S10) one can see how, even though the $R^2$ often hovers around 0, the Spearman correlation reveals the ability of the models to rank the peptides despite the low $R^2$.

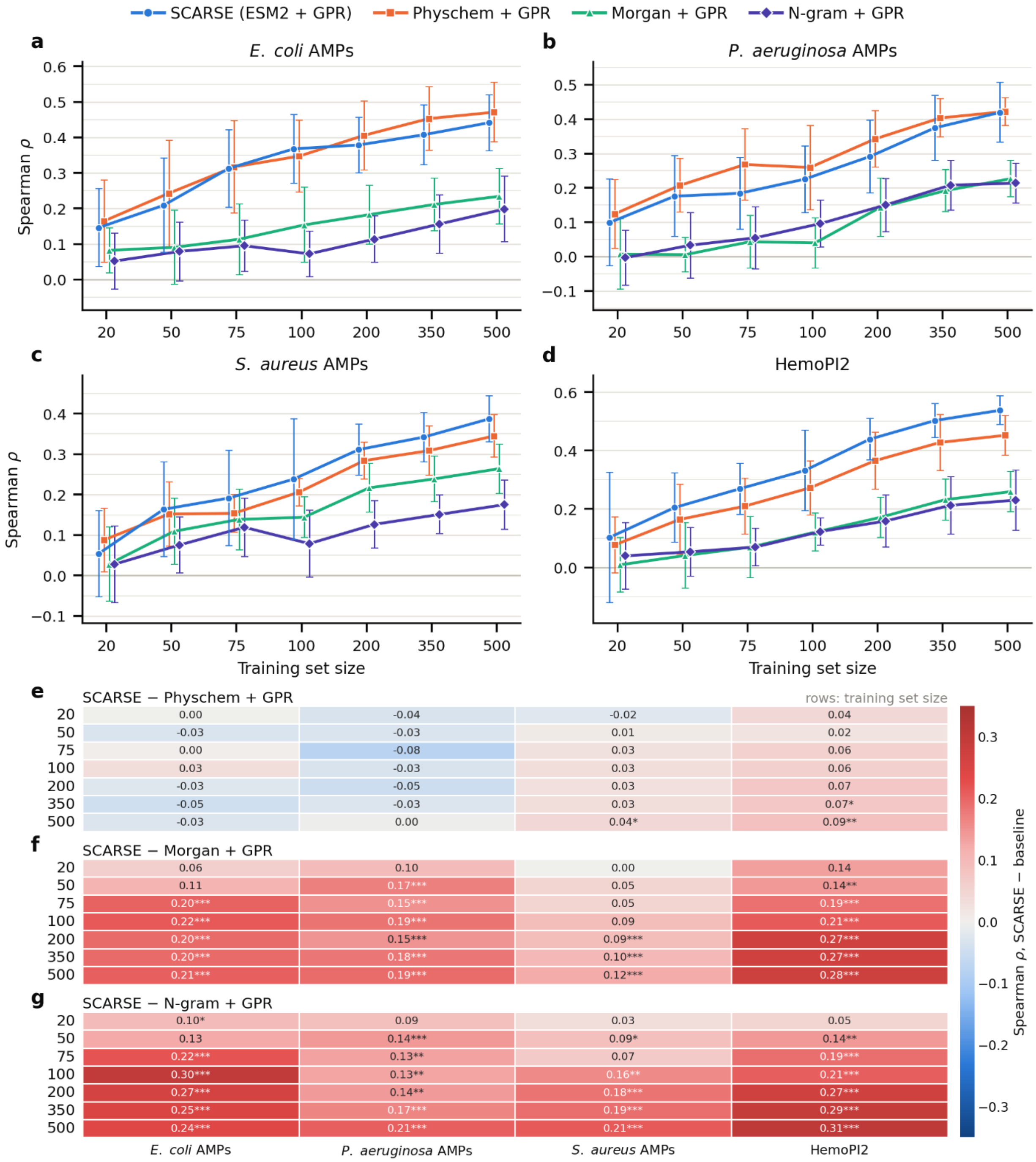


Fig. S10. Benchmarking performance of the peptide datasets (Spearman ρ). (a–d) Benchmarking performance of the *E. coli* AMPs, HemoPI2, *P. aeruginosa* AMPs, and *S. aureus* AMPs datasets in terms of mean Spearman ρ ± std across 10 random seeds. (e–g) Effect-size difference between the means of SCARSE and each baseline together with the Tukey HSD post-hoc statistical test (*p ≤ 0.05, **p ≤ 0.01, ***p ≤ 0.001).

We also report the Top 20 accuracy for the peptide datasets (Fig. S11), analogous to the substitution analysis in Fig. S7.

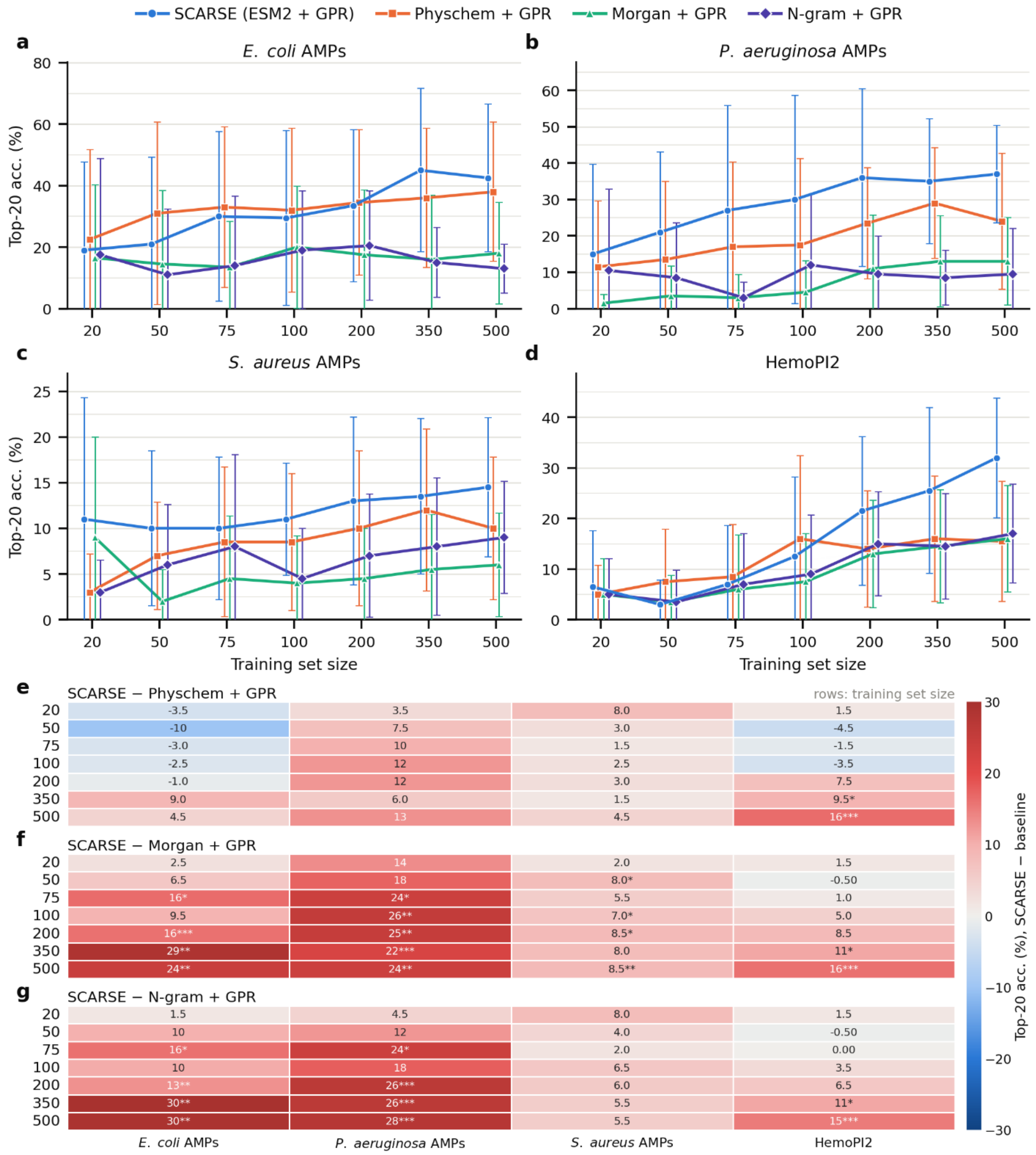


Fig. S11. High-performing selection performance for the peptide datasets. (a–d) Top 20 accuracy (mean ± std across 10 random seeds) for the *E. coli* AMPs, HemoPI2, *P. aeruginosa* AMPs, and *S. aureus* AMPs datasets. (e–g) Effect-size difference between the means of SCARSE and each baseline together with the Tukey HSD post-hoc statistical test (*p ≤ 0.05, **p ≤ 0.01, ***p ≤ 0.001).

From Fig. S9 and Fig. S10 we conclude that these four datasets are generally challenging to predict on using small training sizes, while also highlighting the potential of using ML to predict the ranking of these uncharacterized peptides, as indicated by the Spearman correlation and top 20 accuracy (Fig. S11).

#### Section S5. Active learning methods

We defined two metrics of performance for results in Section S6 (Fig. S12, S13). The normalized target mean per round evaluates the quality of selected peptides relative to the candidate pool and the optimal selection:

$$\text{Target Mean (Round)} = \frac{\mu_{\text{selected}} - \mu_{\text{all,round}}}{\mu_{\text{best,round}} - \mu_{\text{all,round}}}$$

where $\mu_{\text{selected}}$ is the mean true score of the $k$ selected peptides by the model at a given round, $\mu_{\text{all,round}}$ is the mean score of the candidate pool, and $\mu_{\text{best,round}}$ is the mean score of the top $k$ peptides with the highest true scores in the available pool at the given round. Values of 0 and 1 correspond to random and optimal selection, respectively.

To assess accumulated performance, we computed a cumulative normalized target mean:

$$\text{Target Mean (Cumulative)} = \frac{\mu_{\text{current}} - \mu_{\text{all}}}{\mu_{\text{best}} - \mu_{\text{all}}}$$

where $\mu_{\text{current}}$ is the mean score of all peptides currently included in the training set, $\mu_{\text{all}}$ is the mean score across all peptides in the dataset, and $\mu_{\text{best}}$ is the mean score of the top $n$ peptides with the highest true scores in the dataset, where $n$ corresponds to the current size of the training set.

#### Section S6. Active learning simulation performance

This section provides information regarding the selection of sequences throughout the workflow simulations. Fig. S12a-n reports the per-round performance of SCARSE in an active learning workflow scenario for all substitution datasets and the AMP and HemoPI2 datasets, for both the greedy and mixed acquisition strategies. We display the Target Mean (Round) value of the test set across 10 random seeds together with the std, see Section S5 for how these values were calculated.

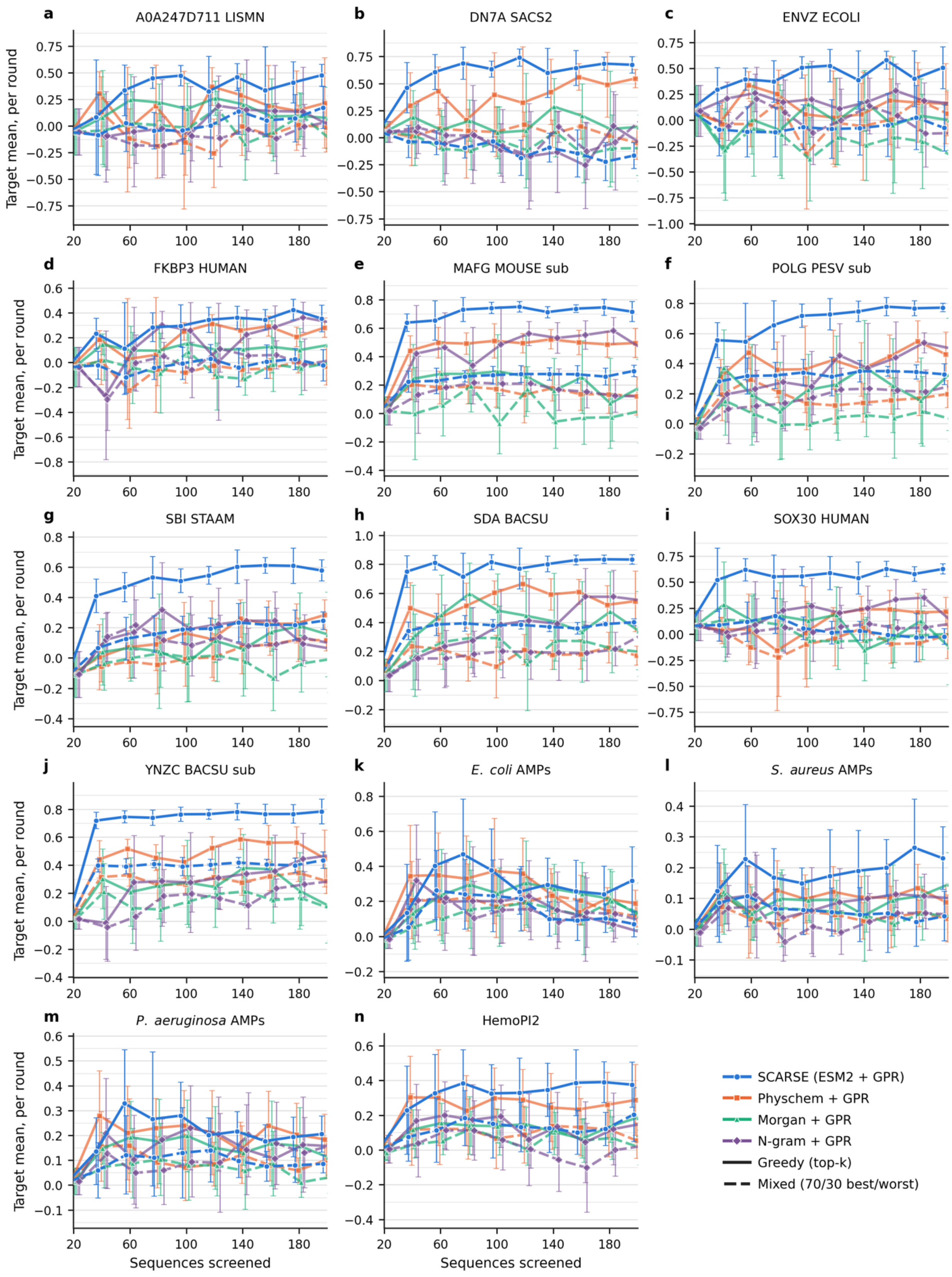


Fig. S12. Per-round workflow simulation performance using SCARSE and the three baselines. (a-n) Performance is reported on the substitution datasets together with the AMP and HemoPI2 datasets, measured as Target Mean

(Round), across iterative selection rounds. Solid lines show greedy top-k selection and dashed lines the mixed (70% best / 30% worst) strategy. Results are displayed as the means ± std across 10 random seeds.

We can see how the performance per round is in general above 0, showing the superior performance of AI-based selection over random sampling, even at very early stages of the discovery workflow. In addition to investigating the candidate selection per round, it would also be of interest to see the cumulative performance of all peptides SCARSE has selected up to a given round (Fig. S13a-n).

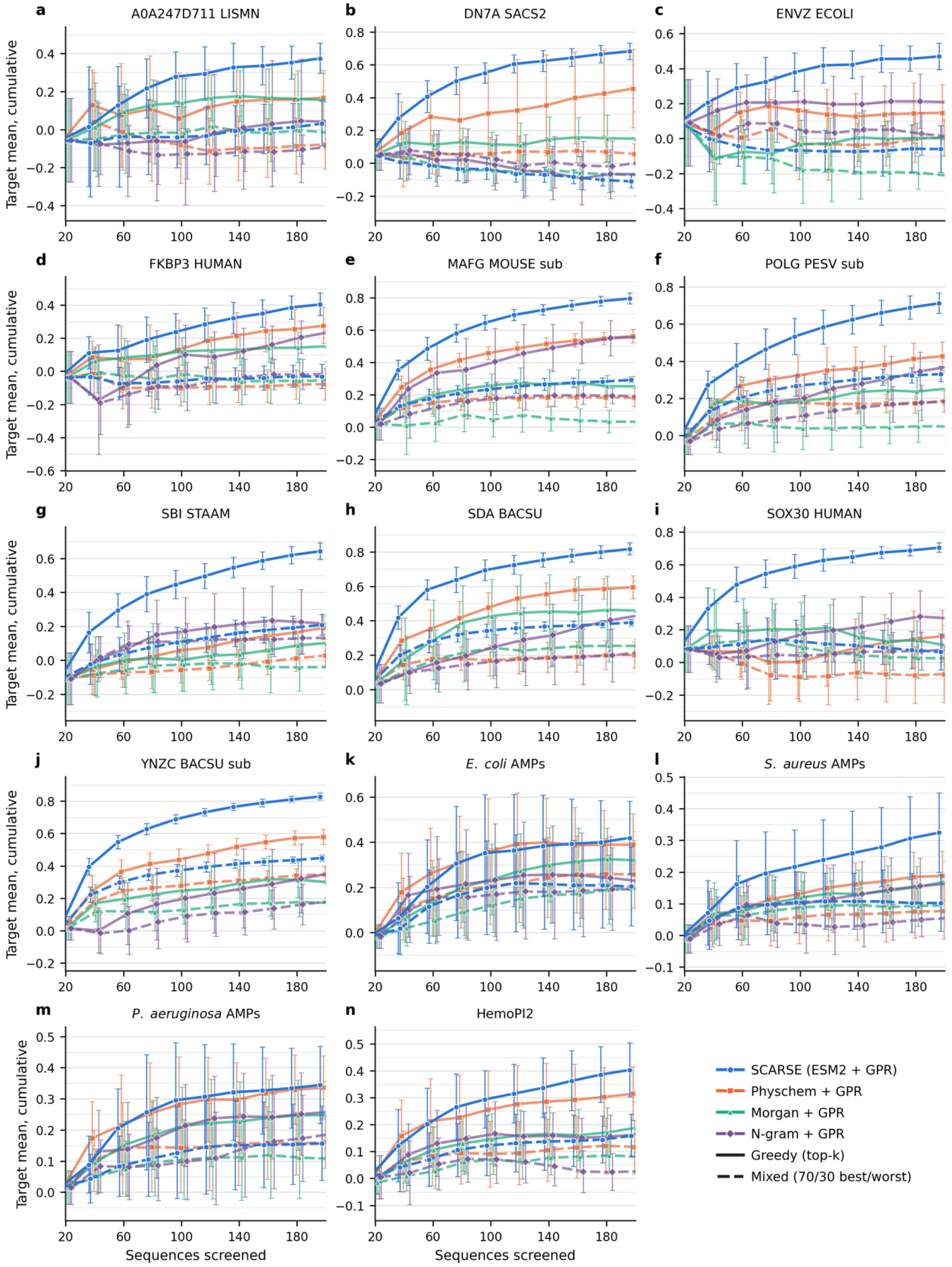


Fig. S13. Cumulative workflow simulation performance across peptide datasets using SCARSE and the three baselines. (a–n) Performance on the substitution datasets together with the AMP and HemoPI2 datasets, measured as

Target Mean (Cumulative), across iterative selection rounds. Solid lines show greedy top-k selection and dashed lines the mixed (70% best / 30% worst) strategy. Results are displayed as the means ± std across 10 random seeds.

The group of peptides selected by SCARSE tend to have a growing performance as the number of peptides screened increases. We note that even low levels of generalizable performance (Fig. S5, S8, S9) are enough for SCARSE to aid discovery in an active learning workflow setting (Fig. S12, S13).

Fig. S14 illustrates the percentage of the actual top 10% best peptide candidates that SCARSE has accumulated at a given number of screened peptides.

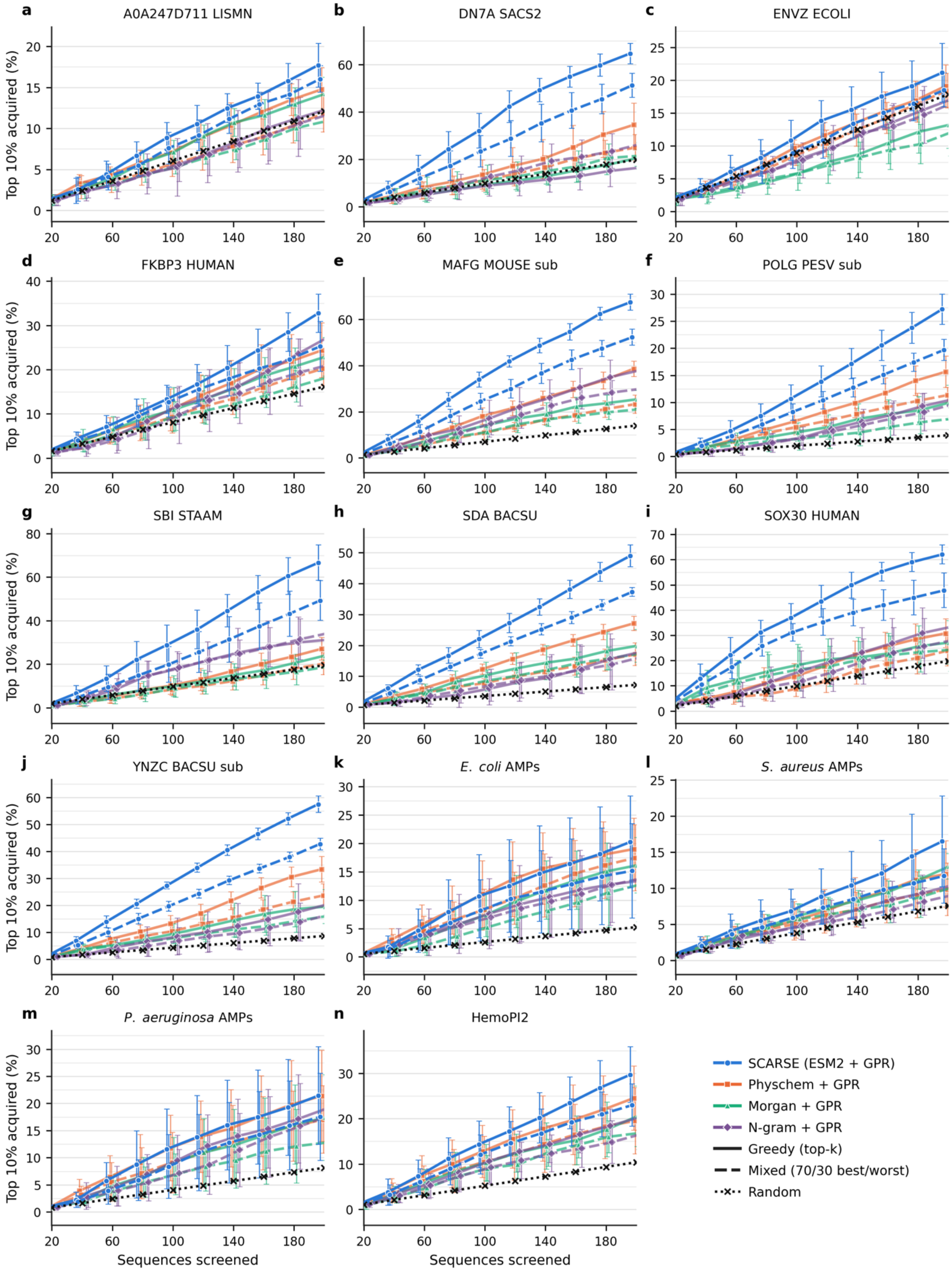


Fig. S14. Active learning high-performing peptide selection performance. (a–n) Percentage of the actual top 10% of peptides acquired by SCARSE and the three baselines up to each selection round. Performance on the substitution datasets together with the AMP and HemoPI2 datasets is investigated. Solid lines show greedy top-k selection, dashed lines the mixed (70% best / 30% worst) strategy, and the dotted line random sampling. Results are displayed as the means ± std across 10 random seeds. The red dotted line indicates mean performance if one were to do random sampling.

This provides clarity into whether SCARSE is capable of detecting high-value peptides in active learning simulations.

The active-learning results in Fig. S12 to S14 are averaged over 10 random seeds. The seeds differ in which 20 peptides form the initial training set and in the stochastic parts of the workflow. For most datasets this averaging is enough and the seed-to-seed variance is modest. The antimicrobial and hemolytic datasets are the exception and show a noticeably wider spread of final active learning outcomes (Fig. S15a). Following individual seeds over the rounds shows that this spread is not present at the start but opens up as screening proceeds (Fig. S15b and S15c). Early greedy choices steer each run toward a different part of the pool and small differences early on are amplified over later rounds.

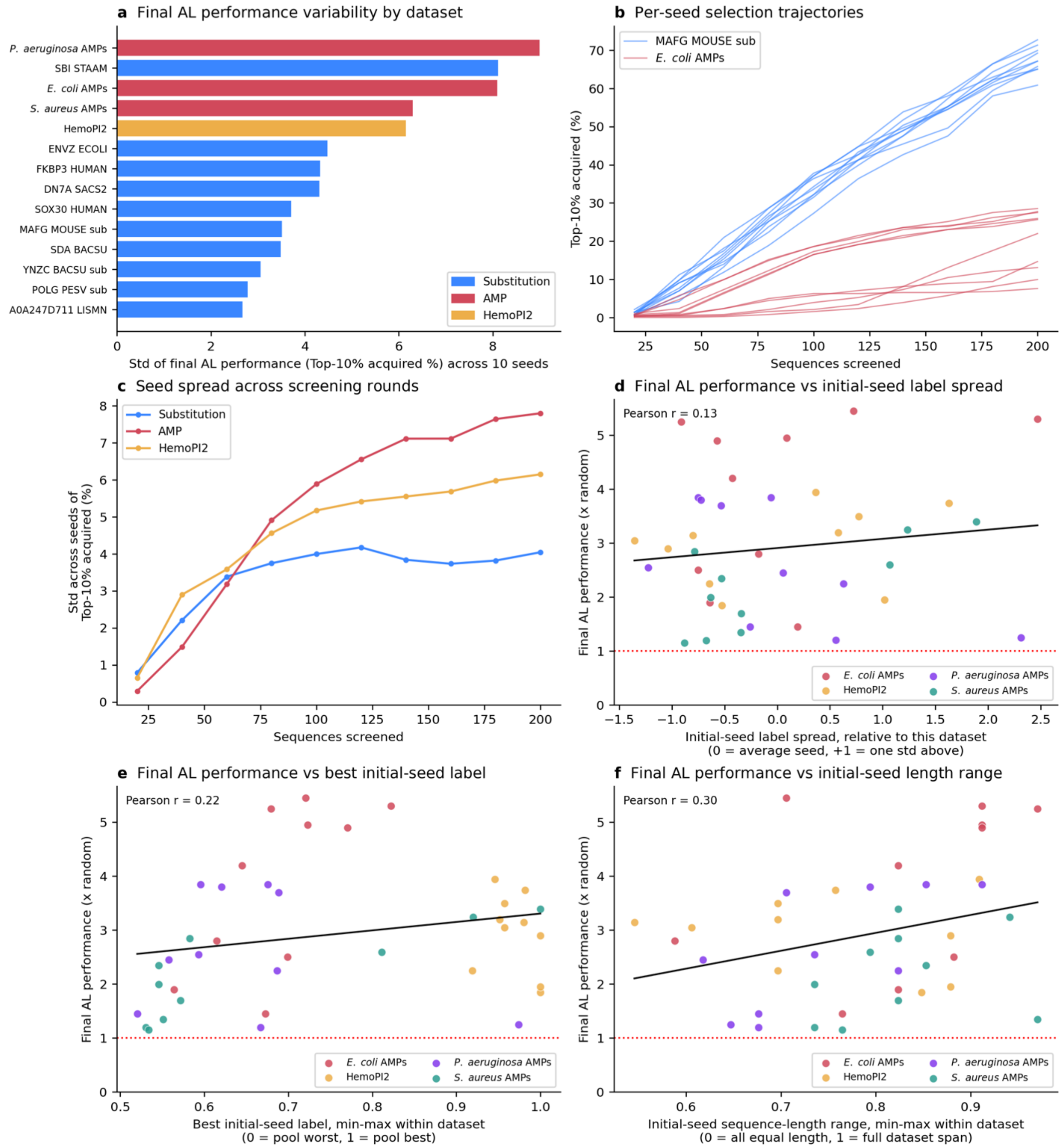


Fig. S15. Seed-to-seed variance of SCARSE greedy selection and its relation to the initial training set. All panels use SCARSE (ESM2 650M + GPR) with greedy top-k acquisition. (a) Standard deviation of the final active learning top 10% acquired performance across the 10 random seeds for each dataset. The antimicrobial and hemolytic datasets sit at the top and most of the substitution datasets at the bottom. (b) Per-seed selection trajectories for one representative substitution dataset (MAFG MOUSE sub) and one antimicrobial dataset (*E. coli* AMPs), where each line is one seed. The antimicrobial trajectories fan out much more strongly. (c) Standard deviation across seeds of the top 10% acquired as a function of the number of sequences screened, averaged within each dataset family. The spread grows over the rounds for every family and remains largest for the antimicrobial datasets, while it flattens out for the substitution datasets after 100 sequences screened. (d–f) Final active learning top 10% acquired performance of each seed, expressed as a multiple of random screening, plotted against three properties of that seed's initial 20-peptide training set: (d) the spread of the initial labels, (e) the best (most desirable) label present in the initial set, and (f) the sequence-length range of the initial set. The three panels use two different normalizations of the x-axis, both applied within each dataset so that datasets on different scales can be shown on a common axis. In (d) the property is standardized across that dataset's ten seeds by subtracting the seed mean and dividing by the seed standard deviation, so a value of 0 marks the dataset's average seed and +1 marks one standard deviation above it; this normalization is relative to the observed seeds, not to the full dataset. In (e) and (f) the property is instead min-max normalized against that dataset's entire possible range: in (e) 0 is the worst and 1 the best peptide present anywhere in the pool, and in (f) 0 means all 20 seed peptides share the same length and 1 means the seed spans the widest sequence-length range the dataset allows. On every x-axis a higher value denotes a more desirable or more diverse initial set, with a lower $pHC_{50}$ (less hemolytic) treated as more desirable for HemoPI2. Points are colored by dataset; the black line is the ordinary least-squares fit and the red dotted line marks random-screening performance. None of the three properties strongly predicts the end point, with the initial-seed length range showing the strongest but still weak association (Pearson r = 0.13, 0.22 and 0.30 for d, e and f respectively).

We asked whether the starting set alone could explain which seeds end up ahead. For each seed we reconstructed its initial 20-peptide training set and summarized it in three ways: the spread of its labels, the single best label it contained, and the range of sequence lengths it spanned. We then compared these to the seed's final active learning performance, measured as the accumulated top 10% of peptides after 200 screened sequences expressed as a multiple of random screening. All three properties show only a weak to moderate association with this outcome: the spread of the initial labels shows almost none (Fig. S15d, Pearson r = 0.13), the best label present in the initial set a weak positive association (Fig. S15e, Pearson r = 0.22), and the initial sequence-length range the strongest but still modest association (Fig. S15f, Pearson r = 0.30). Every seed still finished above random. Since the initial set accounts for little of the spread, the variance more plausibly arises from the round-to-round dynamics of greedy selection rather than from the luck of the initial draw, which remains to be confirmed directly.

In addition, we investigated the correlation between CV $R^2$ values at differently sized random subsets of the substitution, AMP and HemoPI2 datasets in relation to the relative final active learning performance under the greedy top-k selection strategy (Fig. S16). This investigation resulted in a Pearson correlation between 0.8 and 0.83.

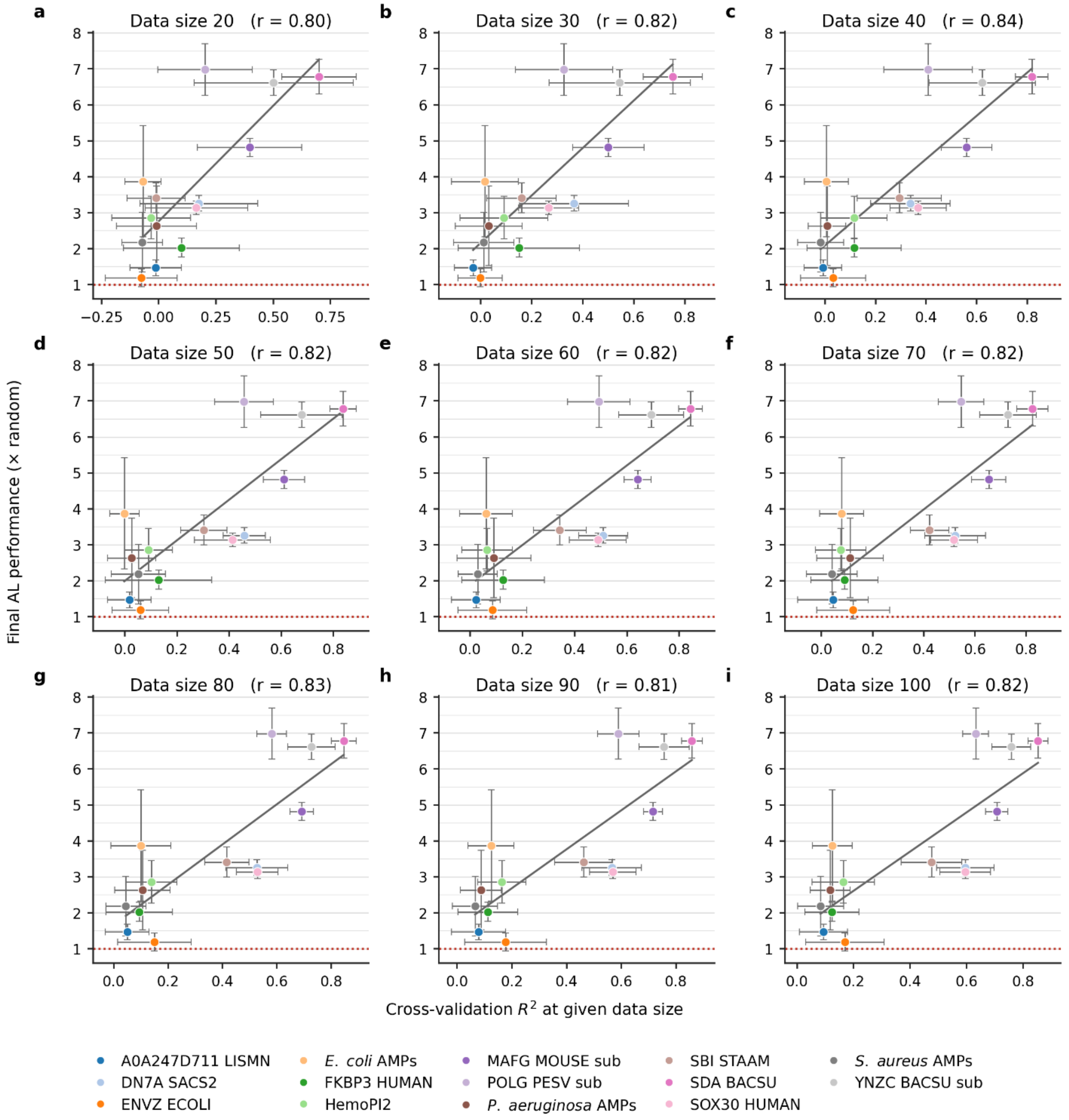


Fig. S16. Final active learning performance correlation analysis. (a-i) Correlation analysis between the final active learning performance and CV $R^2$ score when using SCARSE. The y-axis shows the top 10% acquisition final active learning performance of the active learning workflow simulations in Fig. S14 for SCARSE under the greedy top-k selection strategy, divided by the performance of random sampling, meaning that a value of 1 equals random sampling. The x-axis shows the CV $R^2$ performance for selection of a random subset of data of the size specified in the subplot titles. Mean ± std are calculated based on 10 random seeds both for the CV $R^2$ scores and the final active learning performance. The black line shows a linear regression fitted to the mean performance of all datasets.
